# A genetically-encoded three-colour stress biosensor reveals multimodal response at single cell level and spatiotemporal dynamics of biofilms

**DOI:** 10.1101/2022.09.23.509207

**Authors:** Ahmed E. Zoheir, Morgan S. Sobol, Diana Ordoñez-Rueda, Anne-Kristin Kaster, Christof M. Niemeyer, Kersten S. Rabe

## Abstract

The plethora of chemical, physical, and biological factors that can damage microbial cells has triggered the evolution of sophisticated stress response (SR) mechanisms. While individual SR pathways have been monitored with genetically encoded reporters, sensor concepts for the detection of multimodal effects of stressing conditions in living microorganisms are still lacking. Orthogonally detectable red, green, and blue fluorescent proteins combined in a single vector system, dubbed RGB-S reporter, enable the simultaneous, independent and real-time analysis of the stress response in *Escherichia coli* to physiological stress, genotoxicity, and cytotoxicity. The sensor system can be read out via conventional fluorescence microscopy or microtiter plate analysis and can also be combined with Fluorescent Activated Cell Sorting (FACS) and subsequent transcriptome analysis. Various stressors, such as the biotechnologically relevant 2-propanol, lead to the activation of one, two or all three SRs, which can have a significant impact on non-stress-related metabolic pathways. Implemented in microfluidic cultivation with confocal fluorescence microscopy imaging, the technology enabled spatiotemporal analysis of live biofilms to discover stratified subpopulations of bacteria with heterogeneous stress responses.

## Introduction

The understanding of how microorganisms mediate adaptive changes to ensure survival under changing environmental conditions requires monitoring of the corresponding stress response pathways. RpoS, SOS, and RpoH are critical stress response (SR) pathways that modulate transcriptional pathways through a broad spectrum of stress stimuli, with implications for biofilm formation and proliferation,^1,2^ pathogen virulence,^3^ antibiotic resistance,^4^ evolution^4^ and ecological competition.^5^ RpoS is an alternative sigma factor which regulates, directly and indirectly, about 500 genes in *Escherichia coli* (*E. coli*).^6^ Known as the general stress response, RpoS activation is mainly stimulated by starvation as an indicator of physiological stress.^7^ The SOS response on the other hand comprises more than 50 genes that bear several functions in response to DNA damage induced by chemical, physical or biological agents.^8^ Hence, SOS response upregulation is associated with cell genotoxicity. The alternative sigma factor RpoH is the key regulator of the heat-shock stress response in *E. coli* that encompasses more than 30 genes.^9,10^ Activation of the RpoH response is stimulated by the accumulation of unfolded proteins in the cell as indication of cytotoxicity.

In view of the high relevance of these biological processes for technology and medicine, a comprehensive understanding of the underlying molecular mechanisms is of outstanding importance. For example, monitoring cellular responses to multiple stressors can make an important contribution to understanding cell viability and productivity,^11^ such as product inhibition, nutrient deprivation, pH, or shear stress,^12^ as well as for monitoring a variety of environmental toxicants, such as herbicides or antibiotics.^13^ A multimodal analysis of microbial stress response would therefore be important not only for basic research but also for biotechnological processes.

Towards this goal, several genetically-encoded bacterial biosensors have been developed.^12– 18^ Commonly used reporter elements are the colorimetric β-galactosidase (*lacZ*)^15^ and bioluminescence (*luc, lux*)^16^ reporters. Since these systems usually require cell lysis, multi-step assays, or catalytic reactions which limit online measurement and multi-colour reporting, fluorescence-based reporters have been developed for the analysis of stress response.^17,18^ However, the currently available systems lack the ability to report the multimodal response of living cells with high spatiotemporal resolution. We here describe a genetically-encoded three-colour fluorescent biosensor that simultaneously displays bacterial response to physiological stress, genotoxicity, and cytotoxicity through monitoring of the corresponding SR pathways (Figure 1 and Supplementary Figure 1).

**Figure 1:**
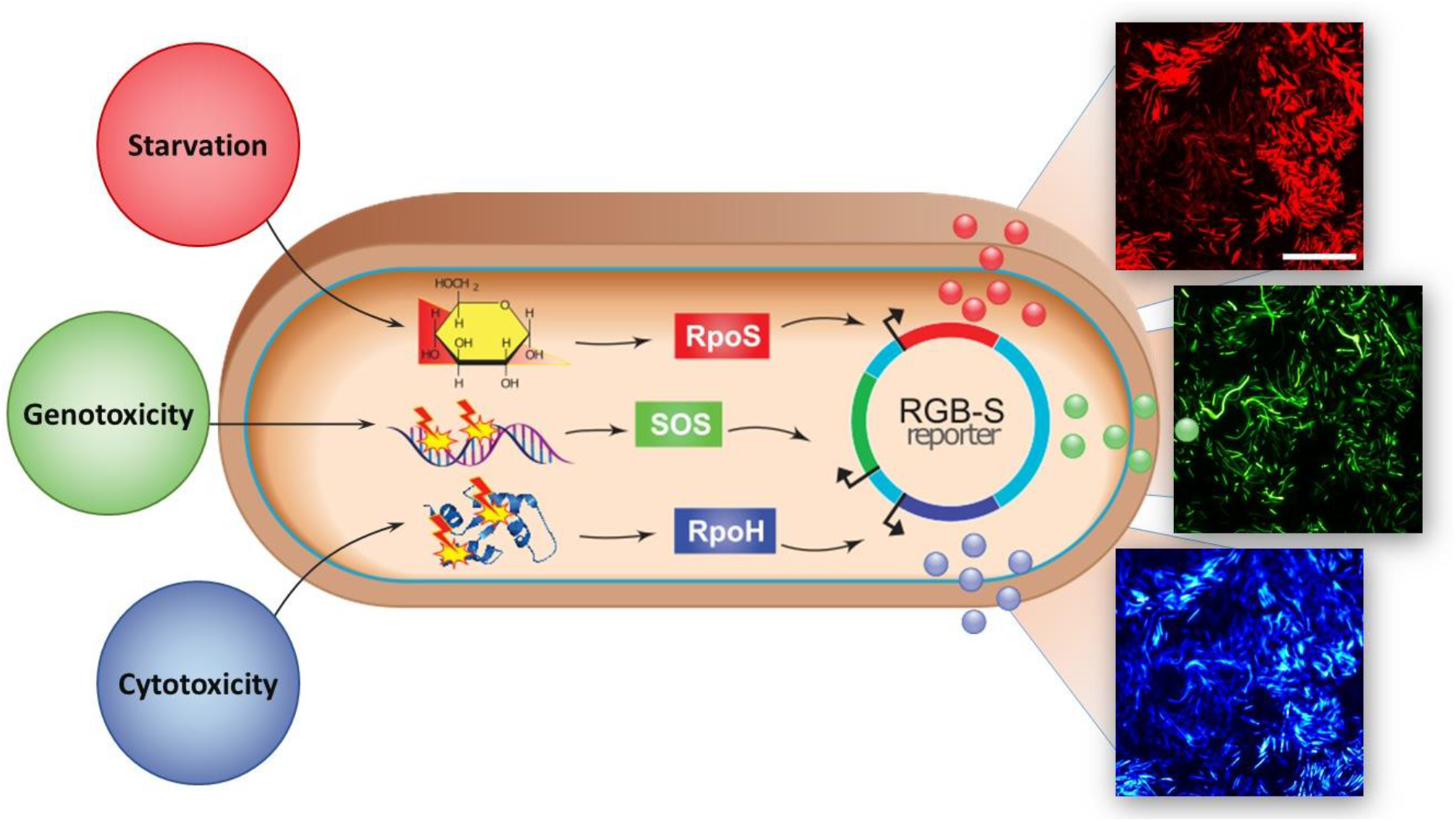
Starvation, genotoxicity, and cytotoxicity are being reported via the RGB-S reporter plasmid in *E. coli*. Activation of the RpoS, SOS, and RpoH stress response pathways by starvation, DNA damaging, or protein misfolding agents leads to expression of red, green, and blue fluorescent proteins, respectively, as indicated by fluorescence microscopy images. Scale bar is 50 μm.

## Results and Discussion

### Sensor construction and validation

To realize a robust multi-colour stress response with high signal-to-noise ratios, three promoters of the model organism *E. coli* K12 MG1566 were selected: P*sulA* is a strongly induced promoter during the SOS response that indicates DNA damage (genotoxicity),^19^ and P*osmY* from the RpoS regulon is an indicator of nutrient starvation, osmotic, and other physiological stresses.^20^ The chaperon promoter P*grpE* is involved in the heat-shock RpoH response, which gets activated due to intracellular accumulation of unfolded proteins that are indicative of cytotoxicity.^16,21^ Three orthogonally detectable fluorescent protein (FP) variants with red (mRFP1),^22^ green (GFPmut3b),^23^ and blue (mTagBFP2)^24^ colours were selected to enable simultaneous read-outs of pathway activation (Supplementary Table 1). The coding sequences for the three FPs were optimized for high protein translation rates^25^ in *E. coli* to secure adequate signals even under conditions of growth-inhibition. Since FP synthesis and maturation play a significant role in fluorescence signal initiation,^26^ we used the enhanced fluorescent proteins GFPmut3b^23^ and mRFP1^22^ which mature faster than the native ancestors. By fusing the FPs downstream of the promoters, the fully assembled RGB-S reporter (for red, green blue stress reporter) contained the genetic elements of P*osmY*::mRFP1 for indication of physiological stress, P*sulA*::GFPmut3b for genotoxicity, and P*grpE*::mTagBFP2 for cytotoxicity (Supplementary Figure 1). To prevent artefacts from translational read-through of one reporter element to the others, strict transcriptional terminators were added in between the sensing elements, resulting in an individually triggered response of the isolated sensor elements (Supplementary Figure 1).

To initially verify the specificity and robustness of the RGB-S reporter in both microscopy analysis and quantitative bulk measurements, its performance was compared with previously published systems. To this end, *E. coli* transformed with the RGB-S reporter was exposed to the herbicide glyphosate. As expected, the upregulation of the starvation response occurred,^27^ resulting in a substantial expression of red fluorescent protein RFP (Figure 2a). Likewise, presence of the antibiotic nalidixic acid (NA) that interferes with the DNA gyrase to impair DNA fidelity and induce SOS response,^28^ triggered the expected expression of green fluorescent protein (GFP). Notably, fluorescence imaging also enabled the identification of unique cell morphologies, such as cellular filamentation. This phenotype is characteristic for late stage SOS response^29^ and it was clearly visible in the GFP channel of cells exposed to NA (Figure 2a) or ciprofloxacin (Supplementary Figure 2). Treatment of the cells with methanol, which is known to affect the membrane permeability besides other cytotoxic modes of action,^30^ led to induction of a strong blue fluorescent protein (BFP) signal (Figure 2a). In addition to microscopy analysis, the response of the RGB-S reporter to stresses could also be monitored in bulk culture to enable detailed quantitative assessment of the kinetics and dose-dependency of drug treatment (Figure 2b and Supplementary Figure 3). Treatment of *E. coli* harbouring RGB-S reporter (RGB-S *E. coli*) with defined dosages of stressors clearly showed the expected dose-dependent increase of the corresponding fluorescent signal over time.

**Figure 2:**
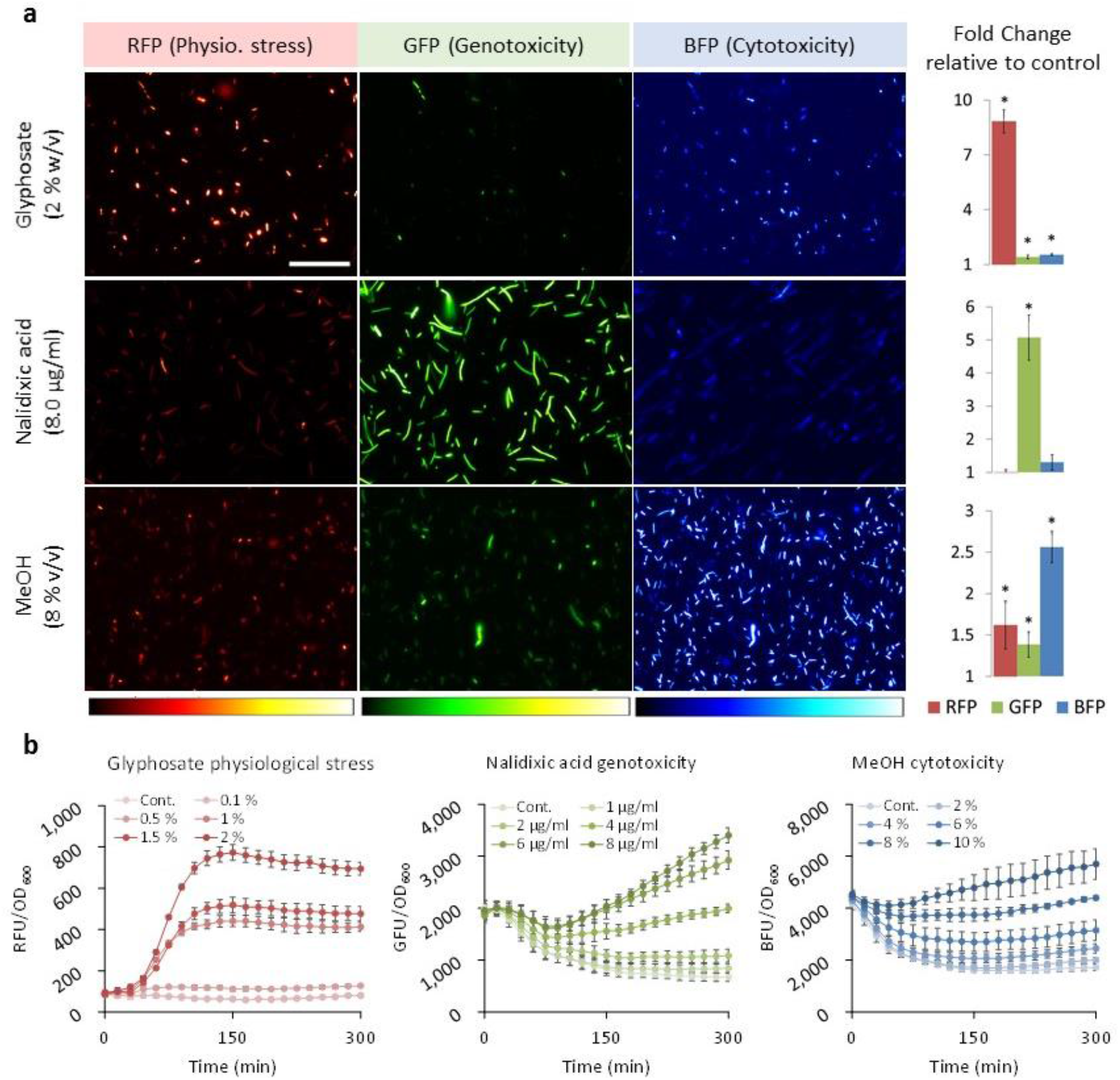
Orthogonal fluorescence readout of stress response reveals time and dose dependencies. **(a)** Fluorescence microscopy images of RGB-S *E. coli* exposed to various stressors for 5 hours. Scale bar is 50 μm. The corresponding quantitative microtiter plate readings, shown as bars, indicate the fold change (FC) of specific fluorescence (FU/OD_600_) over controls that carried the sensor but were not subjected to a stressor. Error bars are the standard deviation (SD) of six biological replicates. (^*^) test and control show significant difference (p≤0.05). **(b)** Time-course measurements of the specific red, green and blue fluorescence response to glyphosate (RFP), nalidixic acid (GFP) and methanol (BFP), respectively, illustrate the dependence of stress response on the stressor dose and exposure time. Error bars are the standard deviation (SD) of six biological replicates.

Furthermore, the RGB-S reporter revealed cellular responses in real-time. For example, the DNA damage induced by NA is cumulative due to the continuous exposure to NA in the medium thus leading to an exponential GFP signal increase over time (Figure 2b). In contrast, a short stimulation by UV irradiation led to an increase of the GFP signal (SOS response due to DNA damage) that reached its peak 45-75 min after the stimulus followed by signal decline likely due to the damage repair or signal dilution by cell growth (Supplementary Figure 3b). Such detailed analyses of the mechanisms of action of cell toxic substances are difficult to achieve with conventional sensor approaches

### Multimodal stress response

The ability to observe responses in real-time revealed variations in signal initiation due to different regulon activation upon employment of variable stressors. For example, we found that cells exposed to glyphosate respond by RFP fluorescence within 15 minutes, whereas exposure to sodium dodecyl sulphate (SDS) led to similar signal intensities only after 4 hours lag time (Supplementary Figure 4).

In addition to correctly identifying the known major response regulon, the RGB-S reporter provided insights into the molecular mechanisms triggered by stressors, which induce more than one response regulon. For example, in case of starvation a bimodal response of RFP and GFP was observed (Figure 3a). While the RFP fluorescence was expected due to the known RpoS induction upon starvation,^32^ the associated 3.5-fold GFP (SOS) response can be attributed to stationary-phase mutagenesis.^33^ Moreover, as a general trend, we observed a triple-modal response for compounds inducing BFP (cytotoxicity), such as ethanol, 2-propanol or acetone (Figure 3a and Supplementary Figure 5). BFP expression is indicative for the accumulation of misfolded proteins, which, in turn, can affect vital functions of cellular components. This explains why RFP and GFP signals are usually associated with the BFP response. On the other hand, the reverse correlation was not observed.

**Figure 3:**
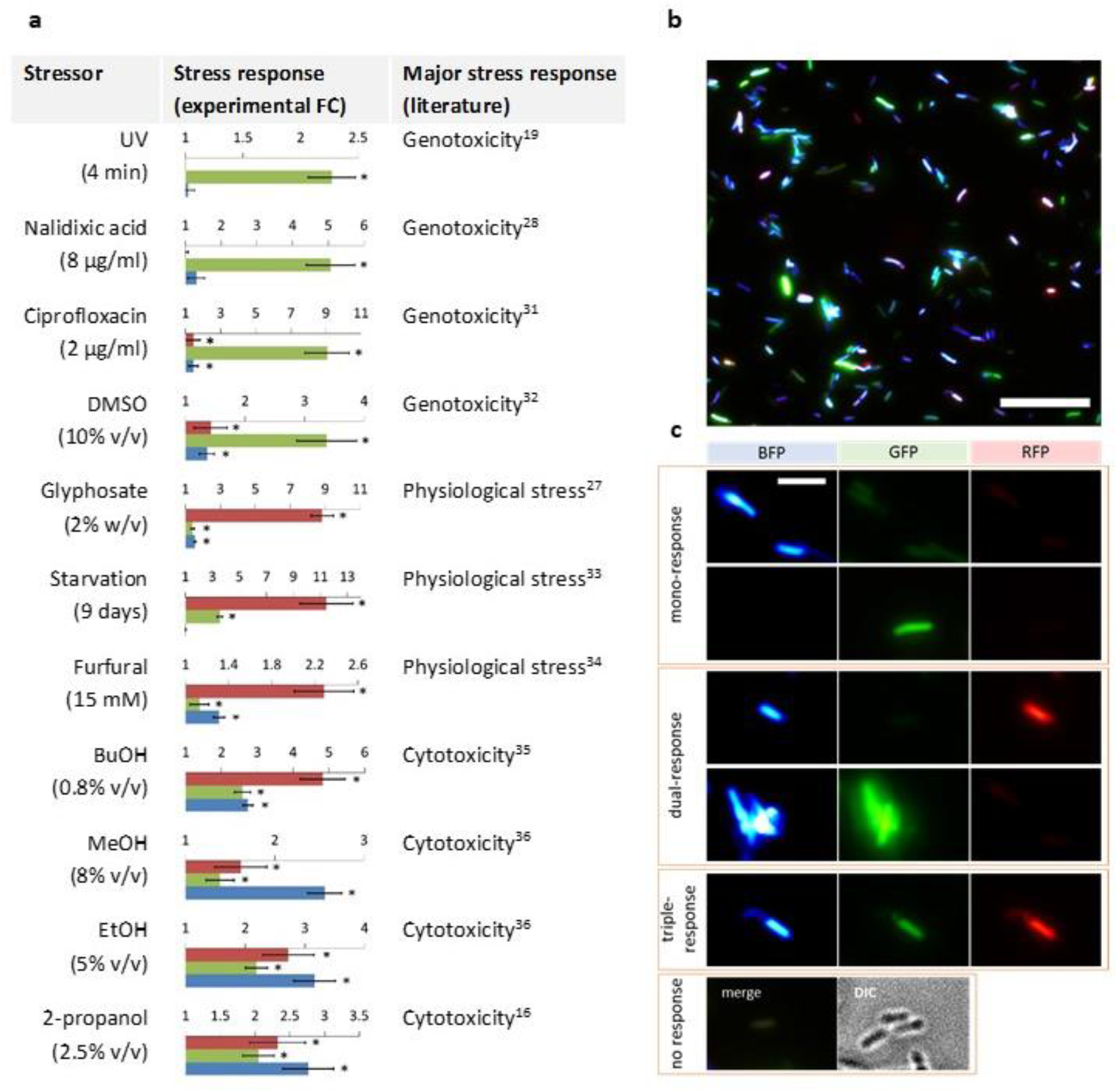
Multimodal effects and heterogenic response to stressors. **(a)** Multimodal response of RGB-S *E. coli* to stress inducers depicted as fold changes (FC) of fluorescence signal relative to non-stressed controls. Error bars are the standard deviation (SD) of six biological replicates. (^*^) test and control samples show statistically significant difference (p≤0.05). Cultures containing the reporter plasmid were incubated for 5 h in Luria-Bertani medium supplemented with kanamycin (LB+kan), except for starvation that was conducted over 9 days. **(b)** Analysis of heterogenic response on the single-cell level to 2-propanol 2% (v/v) determined by microscopy analysis after 5 h incubation. Scale bar is 25 μm. **(c)** Representative images (obtained from **b**) showing cells that display single, multimodal or no stress response. Scale bare is 5 μm.

We then used the RGB-S *E. coli* for direct comparison of mono- and multimodal effects of various stressors (Figure 3 and Supplementary Figure 2 and 5) and found that the RGB-S reporter correctly identified the major modes of action, which had been determined previously by other means (Figure 3a). For example, some antibiotics and UV irradiation were highly specific in inducing GFP fluorescence, thus correlating with their genotoxic potential due to DNA damage (Figure 3a).^19,28,31^ Glyphosate also exhibited a highly specific change in RFP (RpoS) expression due to its inhibition of the shikimate pathway *via* 5-enolpyruvylshikimate-3-phosphate synthase, thus interfering with many metabolic pathways.^27^

The multimodal response capability of the RGB-S reporter was then used for detailed analysis on the single cell level (Figure 3b and c). It was found that the multimodality is based on heterogenic responses of subpopulations, which react on a given stressor in distinctively different ways. For example, after addition of 2-propanol, cells could be assigned to four categories displaying either mono-, dual-, triple-, or no-stress response (Figure 3c). Similar results were obtained from other stressors, such as phenol and butanol (Supplementary Figure 6). While phenotypic heterogeneity is a well described phenomenon attributed to cell-inherent dynamics, transcriptional stochastic effects, and other ecological factors,^34^ the detailed analysis of these processes remains difficult. In this context, the RGB-S reporter can elucidate the molecular basis of such phenomena by enabling the identification and separation of individual subpopulations using established technologies such as fluorescence-activated cell sorting (FACS).

To demonstrate this approach, we sorted approximately 1 million cells of RGB-S *E. coli* treated with 2-propanol 2% (v/v) in two replicates based on the single cell responses into four subpopulations (mono RFP, mono GFP, dual RFP-GFP, and triple RFP-GFP-BFP) using FACS (Supplementary Figures 7). Ttranscriptomic analysis was conducted to determine relative changes in mRNA levels (Figure 4 and Supplementary Figures 8). As expected, the mono and dual responses revealed upregulation in their respective stress regulons. In subpopulations with mono GFP response, the transcription of SOS genes was upregulated, including the *recA* and *lexA* genes, which are known as the main SOS regulators (Figure 4a). Likewise, the RpoS reported gene *osmY* showed predominant transcription both in the mono RFP and dual RFP-GFP subpopulations (Figure 4b). In the dual RFP-GFP subpopulations both the SOS reported gene, *sulA*, as well as the regulator gene, *lexA* showed high transcription levels. These results confirmed the phenotypic analysis of the RGB-S reporter.

**Figure 4:**
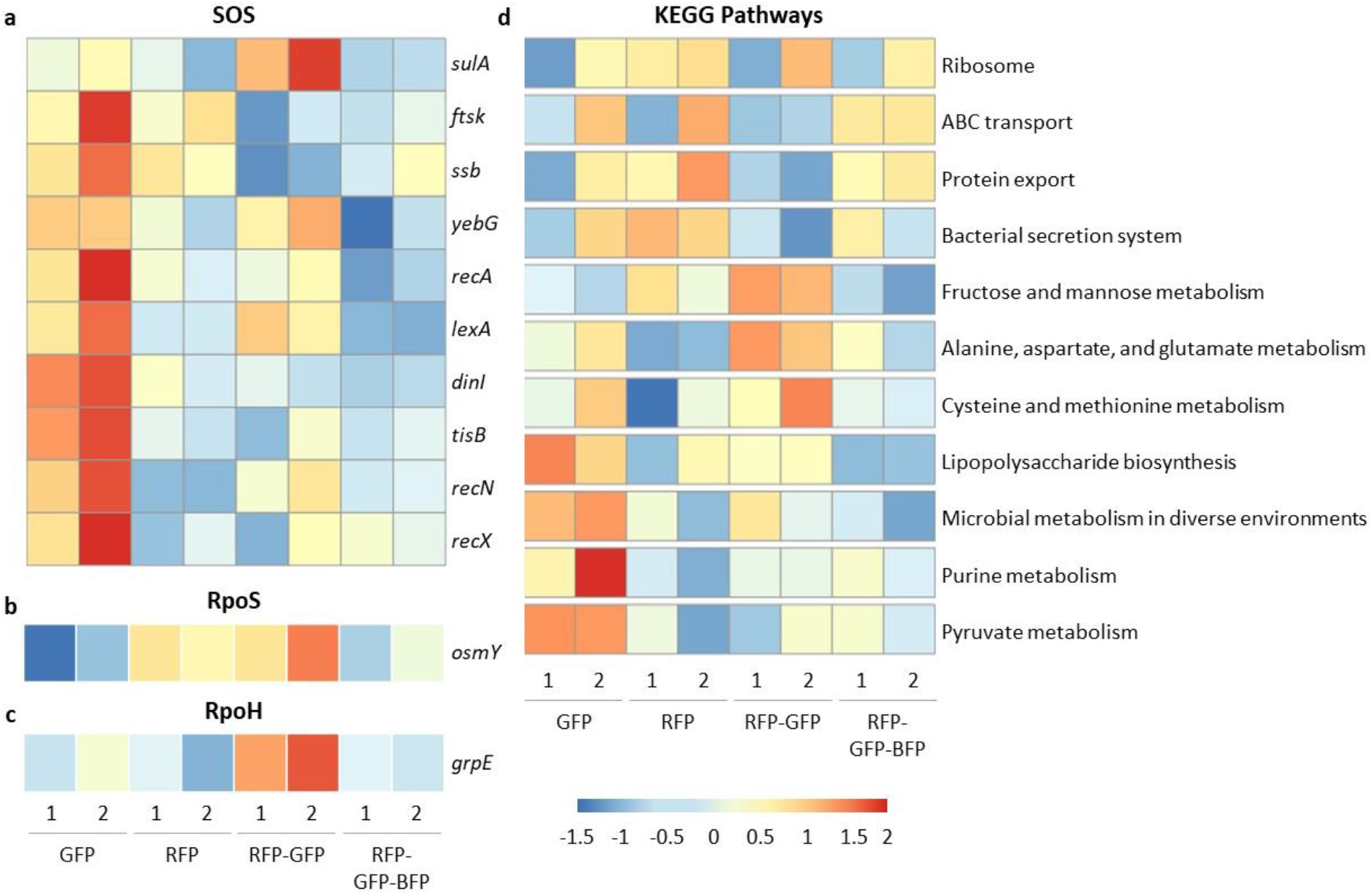
Transcriptomics analysis of SOS, RpoS, and RpoH stress responses and of selected metabolic pathways. RGB-S *E. coli* cells were treated with 2-propanol 2% (v/v) and sorted using FACS. Populations were sorted based on individual cell responses into four subpopulations, mono GFP, mono RFP, dual RFP-GFP, and triple RFP-GFP-BFP, with numbers (1, 2) indicating replicates. Differentially transcribed genes involved in SOS **(a)**, RpoS **(b)**, and RpoH **(c)** stress response pathways were analysed within the four sorted subpopulations. **(d)** Analysis of differentially transcribed metabolic pathways as annotated in the KEGG database of the four sorted subpopulations.

Unexpectedly, however, the RFP-GFP-BFP subpopulations showed downregulation in the transcription of the RpoH reported gene *grpE*, as well as of genes involved in SOS and RpoS responses (Figure 4c). The cells in this population have activated all three stress responses analysed in this study and have started accumulating all three fluorescent reporter proteins, respectively. The intense stress provoked by activating several pathways might, however, lead to the cells shutting down transcription. Such cells would then still show significant fluorescence for all three reporters, but reduced overall transcription levels which would ultimately lead to stopped cell division thus undiluted fluorescent proteins. This hypothesis is supported by the low transcription profile of the RpoH regulon observed in the triple response cells (Supplementary Figure 8). The results suggest that more in-depth time and dose-dependent studies of such subpopulations are needed to elucidate the detailed transcriptional profile of the RpoH regulon to stressors.

Interestingly, transcriptome analysis of the sorted subpopulations revealed other transcriptional profiles altered by the stressor in addition to SR signalling pathways, as shown in particular by enrichment analysis of Kyoto Encyclopedia of Genes and Genomes (KEGG)^35^ metabolic pathways (Figure 4d). For example, lipopolysaccharide biosynthesis, purine, and pyruvate metabolisms were shown to be upregulated in the mono GFP responding populations, while fructose, alanine, and cysteine metabolisms were upregulated in the dual RFP-GFP populations (Figure 4d). Already these first new findings on the modulation of such metabolic pathways clearly demonstrate the enormous potential of the new approach for the identification of unknown stress-induced effects, which could be applicable for the rational design of production strains and further applications in synthetic biology. These novel interactions can further be elucidated *via* systems biology and other computational approaches that utilize transcriptomic data sets from variable stress conditions to unveil the cross-interactions between stress and non-stress related pathways.^36^

### Compartmentalised response in biofilms

Phenotypic heterogeneity is also a distinct feature of bacterial biofilms, which are important in ecology as well as medical and biotechnological settings.^37,38^ In order to investigate living *E. coli* biofilms, we employed a microfluidic flow cell setup^39^ enabling non-disruptive *in situ* analysis of the RGB-S *E. coli* by confocal laser scanning microscopy (CLSM) (Figure 5a, b and Supplementary Figure 9). Imaging analysis of mature biofilms, grown for 96 hours in the absence of stressors, typically revealed signals originating from a few, randomly occurring fluorescent cells at locations in the centre of the microfluidic channel (Figure 5b, “centre”). However, in specific locations, such as at the edge of the fluidic channel where the flow profile leads to accumulated biomass, distinctive patterns of fluorescent cells were visible (Figure 5b, “edge”). For example, a noticeable GFP expression was frequently observed in aggregated cell clusters. Further, an increased RFP expression was often observed in cells near the bottom of the biofilm, presumably indicative for the limitation in nutrients in the respective microenvironments.

**Figure 5:**
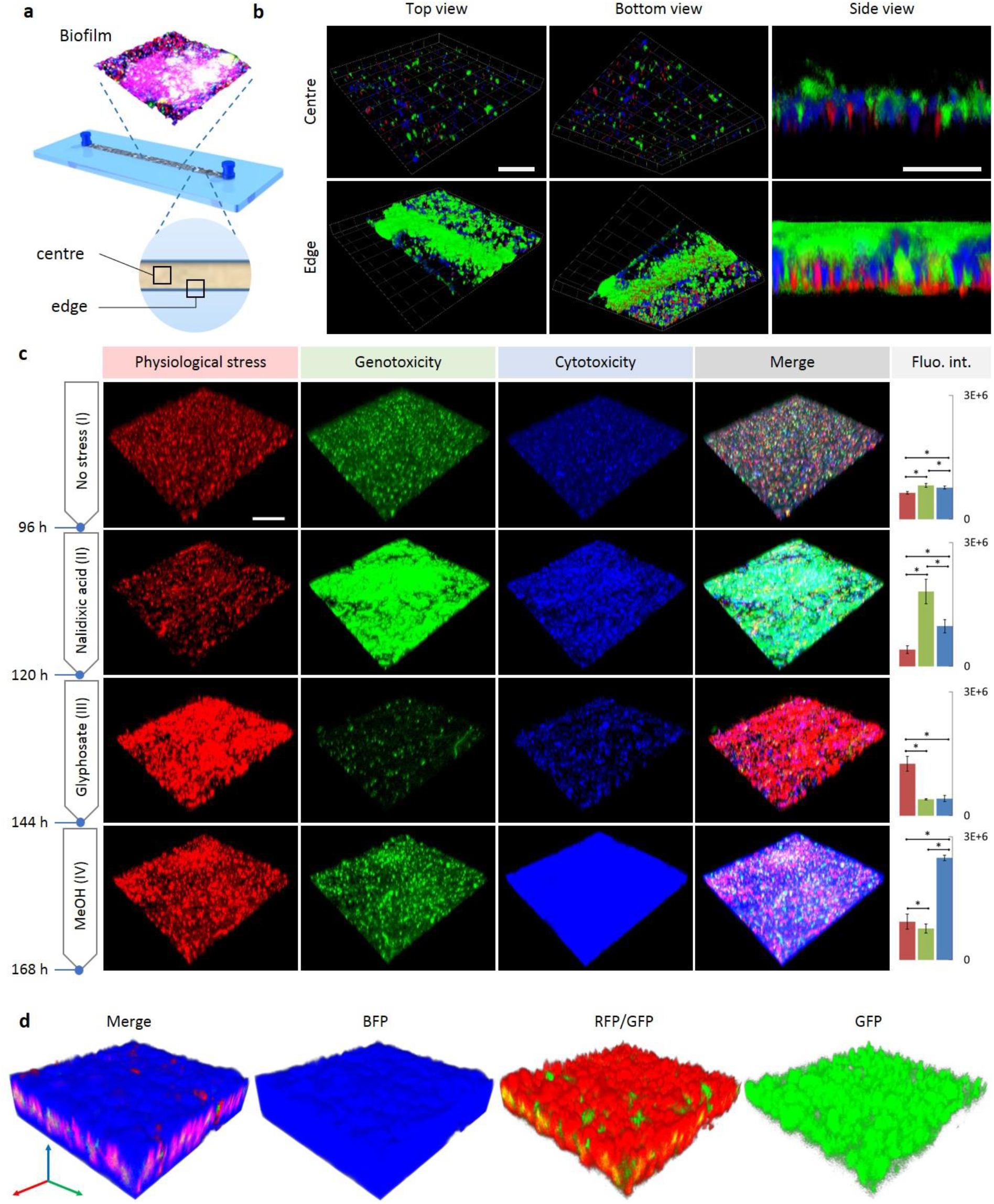
Spatial organization and triggered stress responses in biofilms. **(a)** Schematic illustration of flow channel used for cultivation and real-time imaging of a living RGB-S *E. coli* biofilm grown under continuous flow of 10 μl/min LB+kan. The “centre” and “edge” positions are indicated. **(b)** A mature biofilm grown for 96 h mainly shows a low heterogenic response caused by random occurrence of individual stressed cells (upper row). Near the edge of the flow channel (lower row) GFP-expressing cells dominate the top of the biofilm whereas RFP-expressing cells accumulate at the bottom. Scale bar is 100 μm for top and bottom views, and 25 μm for side views. **(c)** After growth of the biofilm for 96 h only a basal level of stress response can be detected at centre positions (no stress, **I**). After treatment with nalidixic acid for another 24 h (NA, **II**), the cells in the biofilm displayed an elevated GFP signal (genotoxicity). Subsequent administration of glyphosate (Gly, **III**) leads to decreased GFP response with concomitant occurrence of an elevated RFP (physiological stress) response. Further treatment with methanol (MeOH, **IV**) leads to induction of strong BFP signals (cytotoxicity). Scale bar is 100 μm. Quantitative red, green and blue bars (on the right) represent the intensity average of RFP, GFP and BFP, respectively, acquired from nine 100 μm^2^ regions. (*) indicate statistical significance of response means; error bars represent the standard deviation. Additional images of different centre positions are shown in Supplementary Figure 10. **(d)** Internal organization of the three stress responses inside the final MeOH treated biofilm, imaged at a centre position. The biofilm displays a blue fluorescence with a second internal red fluorescent layer and a final inner thin SOS layer. 3D scale bar is 25 μm.

Importantly, the experimental setup enabled the controlled perturbation of biofilms by administration of distinctive stressors, using otherwise unaltered environmental conditions like constant nutritional and flow (Figure 5c and Supplementary Figure 10). For example, after an initial growth phase of 96 hours without stressor (Figure 5c, I), the mature biofilm was sequentially exposed to nalidixic acid (NA), glyphosate (Gly), and methanol (MeOH), respectively, each for 24 hours under otherwise unchanged flow conditions. As expected from the results with the planktonic cultures described above, NA led to a specific increase in the GFP signal (SOS), indicating DNA damage in biofilm-forming cells (Figure 5c, II). Exchange of the genotoxic NA to the physiological stressor Gly not only induced the expected high RFP signal, but also restored the non-stressed GFP state (Figure 5c, III). The same effect was also observed when applying MeOH as the third stressor, which induced a strong BFP response, along with minor RFP and GFP signals (Figure 5c, IV), similar to the multimodal response observed with planktonic cells. The dynamic response of the living biofilm could also be monitored in real-time (Supplementary Figure 11). These results vividly document that biofilm molecular stress-coping mechanisms respond flexibly to environmental changes, which, among other defence strategies, should also contribute to the well-known inherent resilience of biofilms.

Perturbation with stressors also alters the three-dimensional organization in the living biofilm. For example, after 24 hours of MeOH administration, the entire biofilm structure showed a largely homogeneous blue fluorescence, as expected (Figure 5d). However, more detailed analysis with all three colours surprisingly revealed a distinct three-dimensional layered pattern of SRs within the biofilm, indicating inner layers with predominant RFP fluorescence (physiological stress) and a thinner GFP layer (SOS) in the middle. Stress heterogeneity and structural stratification in biofilms have been connected to microgradients of nutrients, oxygen and metabolites.^37,40^ The homogeneous blue fluorescence observed here across the entire biofilm with an average structural thickness of about 50 μm suggests that diffusion of stress factors was not restricted under the conditions used. Therefore, metabolic activity could be the reason for the observed stratification. Since oxygen is rapidly consumed by cell metabolism, the middle red layer (Figure 5d) could result from local oxygen limitation and thus represent the most physiologically stressed and therefore SOS-sensitive zone (green) of the biofilm. Similar effects have been described for *Pseudomonas aeruginosa* biofilms, where oxygen consumption by surface layers of the biofilm could be faster than its diffusion into the lower layers, thus leading to a metabolically active surface in addition to inactive thick bottom layers.^41^ Overall, the observations made here demonstrate for the first time that a living *E. coli* biofilm under continuous flow conditions responds at the cellular level with a dynamic and spatially localized response to stressors. These results underscore the utility of the technology presented here and suggest that a combination with spatiotemporally resolved sampling^39^ and sequencing techniques should enable detailed investigations of the underlying molecular and physiological mechanisms at such complex microbial communities.

In conclusion, the RGB-S reporter described here facilitates multi-stress analyses in bacterial planktonic and biofilm populations, revealing population responses to stresses along with population heterogeneity and spatial organization. Since the novel biosensor is compatible with modern high-throughput screening and imaging systems,^42^ this kind of multi-stress biosensors will enable the acquisition of high-content data to facilitate the fast, robust and comprehensive analysis of bacterial response to transient and permanent environmental changes. We believe that this approach is of great value not only for fundamental research in the context of stress-emergence of antibiotic resistance^43^ and stress heterogeneity within biofilms^37,38^ but also for practical applications in medicine and biotechnology.

## Methods

### Genetic elements

The RGB-S reporter was constructed by fusion of three synthetic sensing elements, cloned into the backbone pMK-RQ [kanR & ColE1 ori] (GeneArt® Gene Synthesis, ThermoFisher Scientific and IDT Inc.) (Supplementary Figure 1). Each sensing construct is comprised of three parts, a) a stress-responsive promoter, b) a fluorescent reporter protein and c) a transcriptional terminator. Based on the main criteria of exhibiting a fast and specific response, stress-responsive promoters were selected based on literature studies. Sequences of the chosen promoters were obtained from the genome sequence of *E. coli* str. K12 substr. MG1655 (GenBank: U00096.3). Fluorescent proteins were chosen with respect to high signal intensity and spectral compatibility. To enhance their translation rates, codons for selected fluorescent proteins were optimized for *E. coli* using the GeneOptimizer™ software (ThermoFisher Scientific).^25^ Sequences of fluorescent proteins as well as transcriptional terminators were obtained from the well documented parts of the Registry of Standard Biological Parts (parts.igem.org). Sequences of all genetic elements are listed in Supplementary Table 2. For handling of genetic designs *in silico*, the software Geneious 9.1.8 (Biomatters Ltd.) was used.

### Cloning

The RGB-S reporter was assembled using the isothermal cloning reaction^44^ Gibson Assembly® Master Mix (NEB inc.) according to the manufacturer’s protocol. If necessary, template DNA was removed by DpnI treatment. 1 μl of methylation-sensitive restriction enzyme DpnI and 2 μl CutSmart® buffer (both from NEB inc.) were added to the assembled reaction and incubated at 37 °C for 30 min. The reaction product was used to transform chemical competent *E. coli* cloning strain DH5α (lab stock) then plated on LB+kan agar and incubated overnight at 37 °C. All plasmids were purified using the ZR Plasmid Miniprep – Classic (Zymo Research inc.) following manufacturer’s protocol, sequence verified (LGC genomics) and stored at -20°C.

The overall cloning strategy is depicted in Supplementary Figure 1. After initial construction of the dual-colour sensor (named RG-S reporter) consisting of P*sulA*::GFPmut3b::terminator_1 and P*osmY*::mRFP1::terminator_2, this vector was linearized using primers O17051-F and O17052-R and assembled with the third sensing element (terminator_3::P*grpE*::mTagBFP2::terminator_4), resulting in the final triple-colour/stress sensing plasmid named RGB-S reporter.

### Stress assay

*Chemicals*. Chemicals used in this study are: EtOH 96% ANALAR, 1-BuOH ANALAR and 2-propanol HPLC grade from VWR international GmbH. Nalidixic acid, ciprofloxacin, dimethyl sulfoxide (DMSO) cell culture grade, chloramphenicol molecular biology grade, isopropyl β-D-1-thiogalactopyranoside (IPTG), kanamycin sulphate, ampicillin sodium salt and glucose monohydrate from PanReac AppliChem. MeOH and acetone were analysis grade, phenol was synthesis grade, acetate 100% and glycerol analysis grade from Merck, Darmstadt, Germany. Glyphosate, Roundup® Gran 420 g/kg as sodium salt from Monsanto, Germany. Sodium dodecyl sulphate (SDS) electrophoresis grade from BioRad. Fructose ≥99% from Sigma-Aldrich. Furfural 99% from Acros Organics.

#### RGB-S E. coli preparation

For stress sensing assays, chemical competent cells of wildtype *E. coli* K12 MG1655 DSMZ 18039 (DSMZ GmbH, Germany) were transformed with the RGB-S reporter (designated as RGB-S *E. coli*) and plated on LB+kan agar as described above. A seedbank of glycerol cryostock of the RGB-S *E. coli* was prepared using a single pure colony to minimize phenotypic heterogeneity in the stress assays. From an overnight LB+kan agar plate, several single colonies were inoculated each into 5 ml LB+kan broth and incubated in 180 rpm shaker at 37°C. From these exponentially growing cultures cryostock aliquots of 15% glycerol were stored at -80°C. A fresh aliquot was used to start every new stress assay experiment.

#### Assay protocol

The assay protocol was designed in order to reliably determine the culture stress state even under high stress levels where no further culture growth was occurring. The stress assay started by using an aliquot of the seedbank to inoculate two independent cultures (Cult. 1 & Cult. 2) each in 5 ml LB+kan broth and incubated overnight in 180 rpm shaker at 37 °C. The next day, cultures were diluted to 1:250 using fresh LB+kan and incubated again at the same conditions for 5-6 hours. After reaching adequate optical density, the cultures were diluted using fresh LB+kan to adjust the OD_600_ to 0.4, which is 2X of the final cell density in the assay plate. At the same time, the chemical stressing agents were diluted in LB+kan to 2X of the final required concentration. The stress treatment started by adding 250 μl diluted RGB-S *E. coli* culture to 250 μl LB-stress mixture (or LB containing no stressor as the control culture) to form 500 μl total volume, which had a final cell density of OD_600_ 0.2 and 1X stressor concentration. Upon mixing, each of the two cultures (Cult. 1 & Cult. 2) were then distributed in three replicates in 96-micowell plate, each well containing 150 μl. In total, six independent biological replicates are analysed for every stress concentration unless otherwise indicated. The microtiter plate was then covered by a fluorescence-compatible transparent film (Lab Logistics Group Inc.) to prevent culture evaporation and incubated in a Synergy H1 microplate reader (BioTek Inc.) with continuous orbital shaking (282 cpm, 3 mm) at 37 °C and measuring the optical density (OD_600_) and the three fluorescence readouts at wavelengths given in Supplementary Table 1 in assigned time intervals.

For UV irradiation, a UV-C germicidal lamp (Philips UV-C, TUV30W G30T8) was used. The lamp was switched-on for 30 minutes before irradiation to reach the maximum power intensity. Two cultures were prepared as stated above and diluted to OD_600_ 0.2 using fresh LB+kan. 1 ml of each diluted culture was poured into 60 mm sterile Petri dish to form a wide area of homogenous thin culture layer and exposed in duplicates to UV light for different periods. Irradiated cultures were then distributed immediately each into triplicate in the 96-microtiter plate and incubated as previously described. Control cultures were treated based on the same procedure but without UV exposure.

#### 2D Fluorescence imaging

For 2D microscopic imaging, an ApoTome inverted fluorescence microscope (Carl Zeiss Inc.) was used. 10 μl of a stressor-treated culture was loaded on a microscope glass slide and covered by a glass slip then imaged using the compatible three light filter sets corresponding to the three fluorescent proteins (Set 43 HE for RFP, Set 44 for GFP and Set 49 for BFP) as indicated in Supplementary Table 1. Before imaging the stressor-treated cultures, the image acquisition settings were set to the blank using a non-stressed control culture. The three fluorescence channels were imaged for the same field by AxioVision software (Carl Zeiss Inc.) and their fluorescence intensity was displayed in a lookup table (LUT) as a linear gradient created by ImageJ^45^ (imagej.nih.gov/ij).

### Fluorescence-activated cell sorting (FACS)

#### Culture preparation and cell sorting

The RGB-S *E. coli* strain was treated with 2-propanol 2% (v/v) for 5h at 37°C in two cultures of 500 mL each, utilizing the standard stress assay protocol described above. Untreated cultures were prepared under the same conditions as the control. Control cells were then diluted with 1X phosphate buffered saline (PBS) to an approximate concentration of 1 × 10^6^ cells mL^-1^ and analyzed in the cell sorter to set the background fluorescence signal threshold for the sorting of the treated samples. Afterwards, populations from RGB-S *E. coli* treated cells were also diluted with PBS and sorted using a MoFlo XDP High-Speed Cell Sorter (Beckman Coulter) equipped with 405 nm, 488 nm, and 561 nm laser lines. Post-acquisition analysis was done with FlowJo software (BD Biosciences). Both non-stressed control sample and subpopulations of stressed mono RFP, mono GFP, dual RFP-GFP, triple RFP-GFP-BFP with stress fluorescence signals higher than the control background threshold were sorted in duplicates. Gating of the aforementioned populations is shown in Supplementary Figure 7. One millions cells were sorted for each sample (except for RFP-2 and RFP-GFP-2 which received 0.7 and 0.75 million cells, respectively) into 250 μL of RNAPure™ peqGOLD (VWR International GmbH), then immediately vortexed and kept on ice until extraction.

#### RNA extraction and library preparation

RNA was extracted with the Direct-zol RNA Miniprep Kit (Zymo Research Inc.), following the standard protocol. DNase treatment was applied after extraction using the TURBO DNA-free™ kit (Invitrogen) and the RNA stored in 1 μL of RNasin® Ribonuclease Inhibitor (Promega Corporation). RNA was quantified with the Qubit™ RNA HS Assay Kit (Invitrogen). Immediately, RNA libraries were prepared using the Zymo-Seq RiboFree® Total RNA Library Kit (Zymo Research Inc.), with some modifications. These included extending the first-strand cDNA synthesis incubation at 48°C to 1.5 hours, keeping the RiboFree® depletion step to 2 hours even when RNA concentrations were < 250 ng, using RNA Clean & Concentrator-5 (Zymo Research Inc.) for the first cleanup after the RiboFree® depletion step (according to Appendix E), and repeating the final bead cleanup step after PCR twice to remove adapter dimers. Library concentration and size was quantified with Qubit™ dsDNA HS Assay Kit (Invitrogen) and Bioanalyzer High Sensitivity DNA kit (Agilent Technologies Inc.). RNA libraries were sequenced using an Illumina NextSeq 550 with the High Output Kit v2.5 -150 Cycles (2 × 75 bp paired-end) (Illumina Inc.).

All RNA extractions and library preparations were done under a laminar flow PCR workbench (STARLAB International GmbH) decontaminated with either RNase AWAY® or DNase AWAY® (Molecular Bio-Products Inc.) and UV.

#### RNA-seq data processing and analysis

The sequence reads were quality checked using FastQC v0.11.9 (www.bioinformatics.babraham.ac.uk/projects/fastqc) and quality-trimmed using Trim Galore.^46^ The rRNA reads were removed using SortMeRNA v4.3.4 (silva-bac-16s-id90 and silva-bac-23s-id98 databases for reference) with default settings.^47^ The filtered reads were mapped to the *E. coli* MG1655 (ASM584v2) genome, edited to include the sensor construct sequences, with the STAR aligner v2.7.6a.^48^ Intron alignment was disabled by setting --alignIntronMax 1, and read pairs were kept if the length-normalized alignment score and the length-normalized number of matched bases were at least 0.5. Read counts for each gene were determined using the featureCounts.^49^ Counts were normalized and differentially transcribed genes were analysed (padj = < 0.05) with DESeq2 in R v3.6.3.^50,51^ The design for DESeq2 was set to test for variation between the four different RGB-S *E. coli* subpopulations. The Likelihood Ratio Test (LRT) was used as the statistical test. The control samples were used to calculate the log_2_ fold changes between all samples, but were not used for determining significant genes between the four subpopulations. Significantly transcribed genes were then used as input for Gene Set Variation Analysis (GSVA) to determine enriched Kyoto Encyclopedia of Genes and Genomes (KEGG) pathways.^35,52^ Here, the number of minimum required genes was set to 5.

### Microfluidic biofilm growth and CLSM imaging

For 3D biofilm imaging, the RGB-S *E. coli* strain was grown in a straight flow-cell microfluidic chip. The gas-permeable microfluidic structure was made of elastomeric polydimethylsiloxane (PDMS) (Sylgard 184, Down Corning) and bonded to a Cyclo Olefin Polymer (COP) film (HJ-Bioanalytik GmbH, Germany) (Supplementary Figure 9). The biofilm was grown in the chip mounted on the stage of a confocal laser scanning microscopy (CLSM) (ZEISS LSM 880, Carl Zeiss Inc.) under a constant temperature of 37 °C and a continuous LB+kan flow of 10 μl/min using a Nexus 3000 syringe pump (Chemyx Inc.). The initial mature biofilm was grown for 96 h. After biofilm maturation, image acquisition settings were adjusted in order to visualize each basal stress response according to the control biofilm before applying the stressors. After this initial setting, the acquisition parameters were kept identical throughout the entire experiment. LB+kan with SOS inducer nalidixic acid 6 μg/ml was pumped through the chip at the same flow rate for 24 h, followed by LB+kan + glyphosate 2% (w/v) for 24 h, and lastly followed by methanol 8% (v/v) for 24 h. Images were acquired using the software ZEN 2.0 black (Carl Zeiss Inc.) with the fluorescence acquisition spectra shown in the Supplementary Table 1. 3D biofilm images were constructed using the software ZEN 2.3 Blue Edition (Carl Zeiss Inc.).

To generate the fluorescence intensity bars shown in Figure 5c and Supplementary Figure 10, an orthogonal *xy* 2D image summing the *Z*-stack layers was generated by ZEN 2.3 Blue Edition for a total of 9 field-of-view positions. Each field-of-view was a square of 100 μm^2^. These regions were used to calculate the integrated fluorescence intensity (IntDen) for the three channels using ImageJ.

### Statistics

Microsoft Excel was used for data analysis. IBM SPSS Statistics v.24.0 (IBM Inc.) was used for statistical significances determination. When applicable, Kolmogorov-Smirnov and Levene tests were used for normality and homogeneity determination, respectively. Parametric 3-colour groups were compared by One-way ANOVA followed by LSD as a post hoc analysis, while nonparametric groups were analysed by Kruskal-Wallis Test. Significance of independent normal means were determined by Independent Samples T-Test, and not-normal means by Mann-Whitney U Test. Significance of dependent normal means were determined by Paired Sample T-Test, and not-normal means by Wilcoxon Signed Ranks Test. All tests were done at 95% confidence using six replicates, unless otherwise indicated.

## Data availability

All genetic information used or generated throughout this study were made available in the Supplementary Table 2. RNA-seq data was submitted to the National Centre for Biotechnology Information’s (NCBI) Gene Expression Omnibus (GEO) under accession number GSE211360. The materials and data that support the findings of this study are available from the corresponding author upon reasonable request.

## Acknowledgements

This work was supported through the Helmholtz program “Materials Systems Engineering” under the topic “Adaptive and Bioinstructive Materials Systems”. A.E.Z. thanks the German Academic Exchange Service (DAAD) and the Egyptian Ministry of Higher Education (MOHE) for funding his PhD scholarship. We thank Tim Scharnweber, Minja Celikic, and Yong Hu for help on CLSM imaging.

## Author contributions

A.E.Z., C.M.N and K.S.R. conceptualized the research. A.E.Z. conducted all genetic engineering, stress assay and biofilm experiments and analysed all respective data. D.O.R. and A.E.Z. performed the FACS analysis. M.S.S. performed the transcriptomic analysis. A.E.Z., M.S.S, D.O., A.K.K., C.M.N. and K.S.R wrote the paper.

## Competing interests

The authors declare no competing interests.

## Notes

### Competing Interest Statement

The authors have declared no competing interest.

