## Supplementary Information for "A genetically-encoded three-colour stress biosensor reveals multimodal response at single cell level and spatiotemporal dynamics of biofilms"


| Content | Page No. |
| --- | --- |
| Supplementary Figure 1 | 2 |
| Supplementary Figure 2 | 3-4 |
| Supplementary Figure 3 | 5 |
| Supplementary Figure 4 | 6 |
| Supplementary Figure 5 | 7-8 |
| Supplementary Figure 6 | 9 |
| Supplementary Figure 7 | 10 |
| Supplementary Figure 8 | 11 |
| Supplementary Figure 9 | 12 |
| Supplementary Figure 10 | 13 |
| Supplementary Figure 11 | 15 |
| Supplementary Table 1 | 16 |
| Supplementary Table 2 | 17-18 |
| References | 18 |

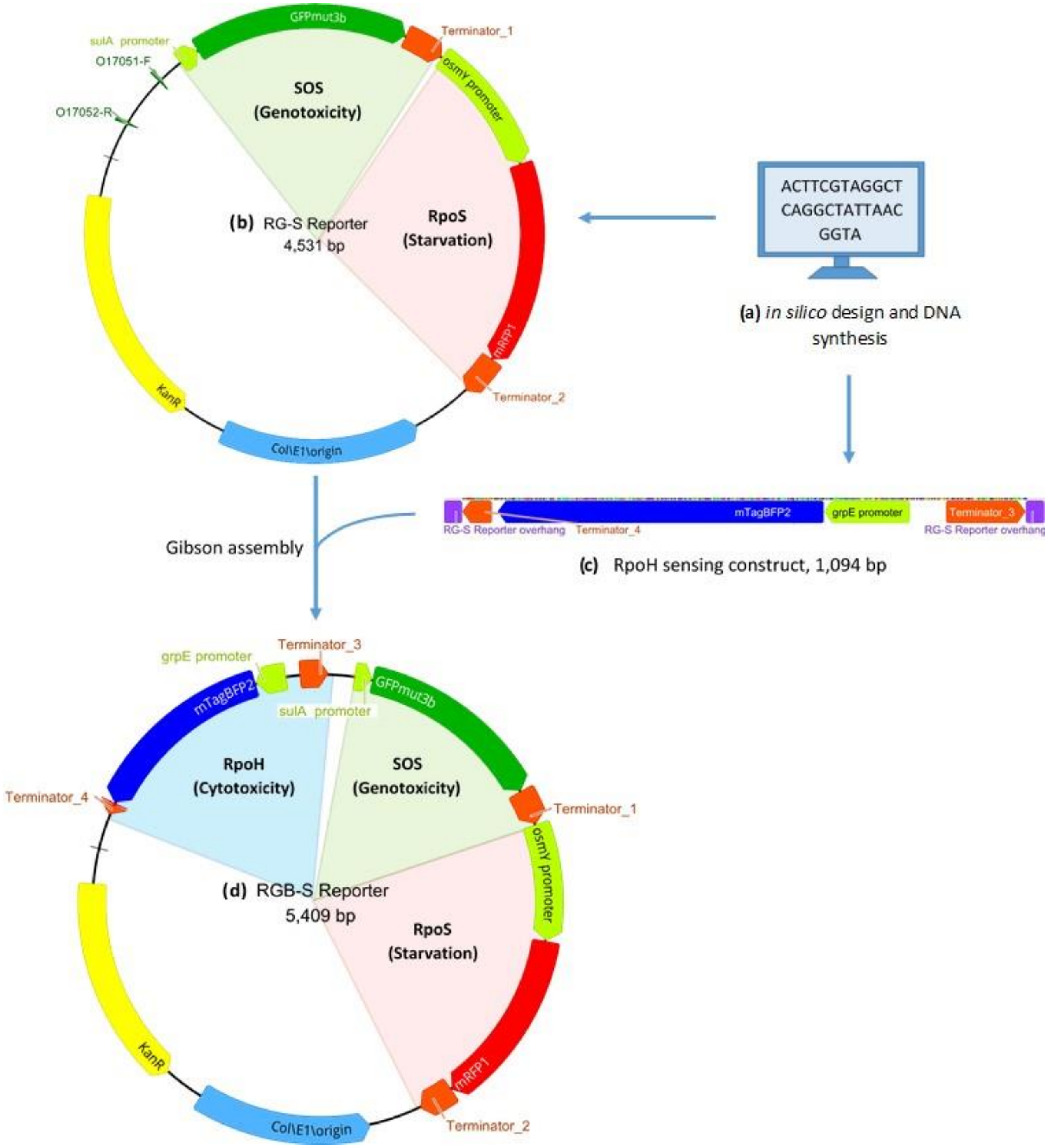


**Supplementary Figure 1**: **Schematic illustration of the design and construction of the RGB-S reporter.** The RGB-S-plasmid was constructed in two steps. **(a, b)** Initially, a dual-colour biosensor RG-S reporter, for sensing SOS and RpoS responses was designed *in silico* and prepared by DNA synthesis (Geneart, Thermo Scientific Inc.). This plasmid was then linearized using the primers O17051-F and O17052-R and the **(c)** synthetic RpoH sensing construct (obtained by *in silico* design and DNA synthesis, IDT Inc.) was added via Gibson isothermal assembly to form **(d)** the final three-colour biosensor, dubbed as RGB-S reporter.

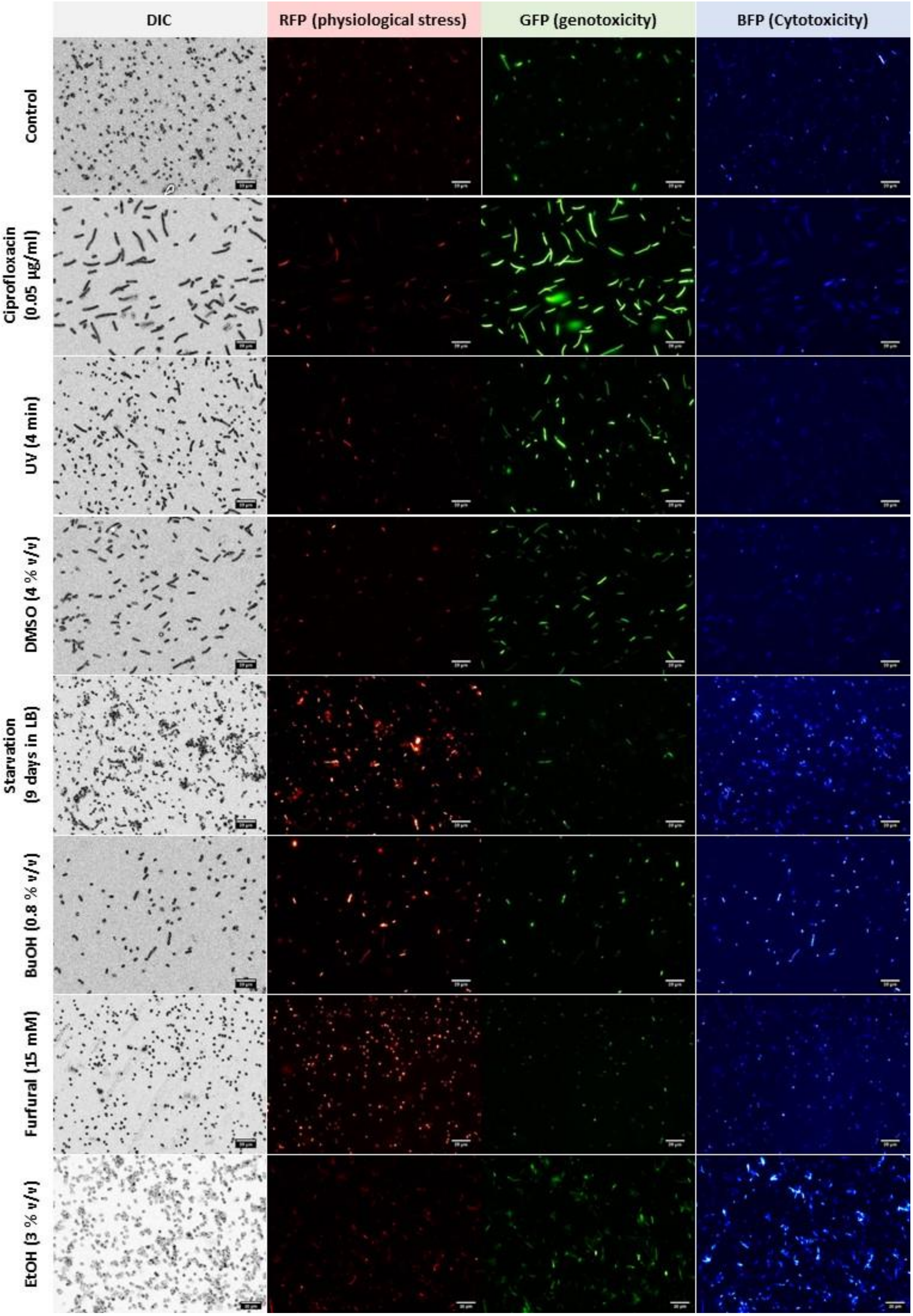
DIC
RFP (physiological stress)
GFP (genotoxicity)
BFP (Cytotoxicity)
Control
Ciprofloxacin (0.05 µg/ml)
UV (4 min)
DMSO (4 % v/v)
Starvation (9 days in LB)
BuOH (0.8 % v/v)
Furfural (15 mM)
EtOH (3 % v/v)

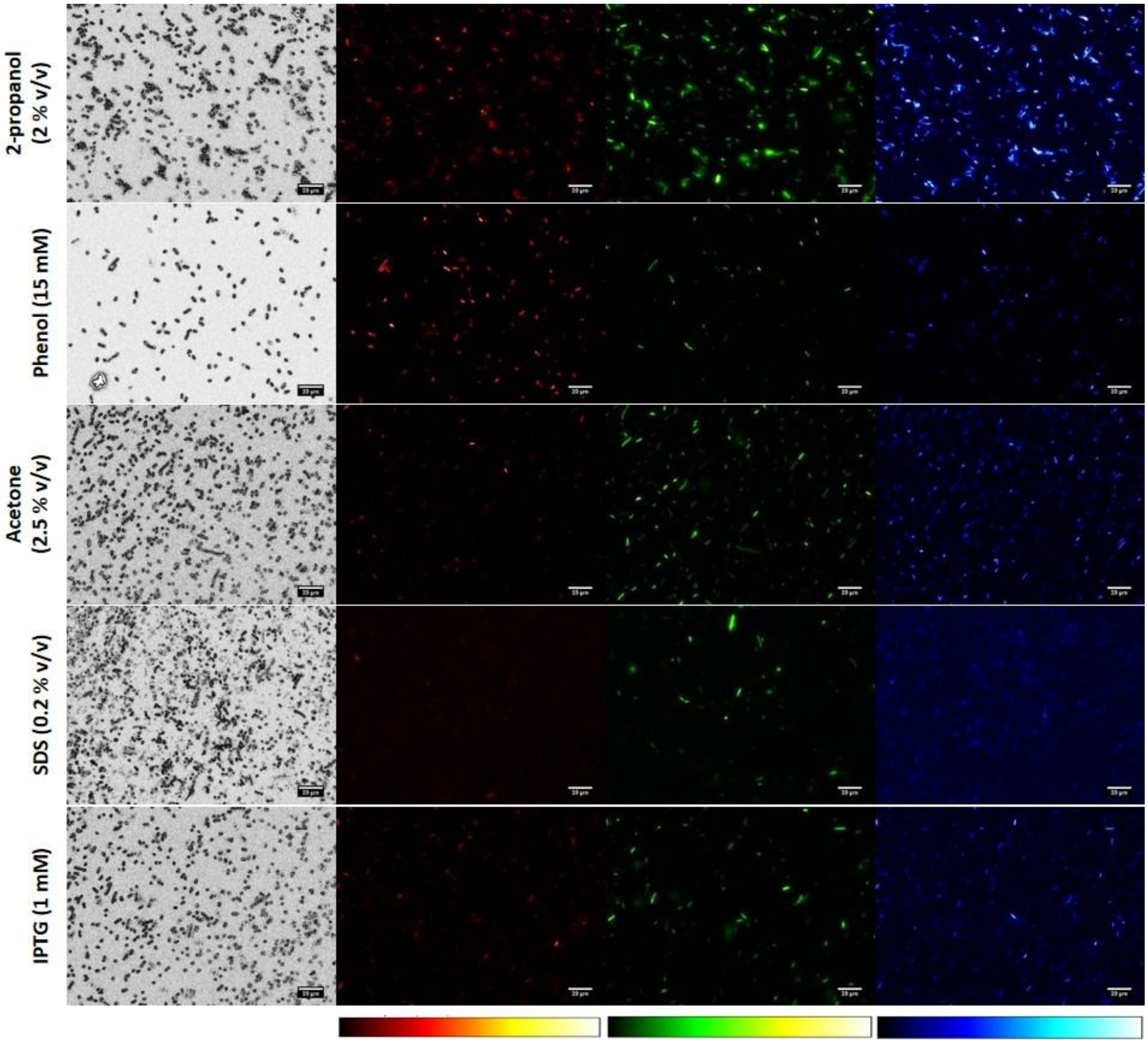


**Supplementary Figure 2: Representative fluorescence images of the RGB-S *E. coli* after treatment with various stressors.** Differential interference contrast (DIC) as well as the three fluorescence channels were taken for the same field of view after 5 h of stress exposure with identical imaging settings for all experiments. Fluorescence intensity is displayed in linear colour gradient created under identical settings using ImageJ. Scale bars are 20 µm.

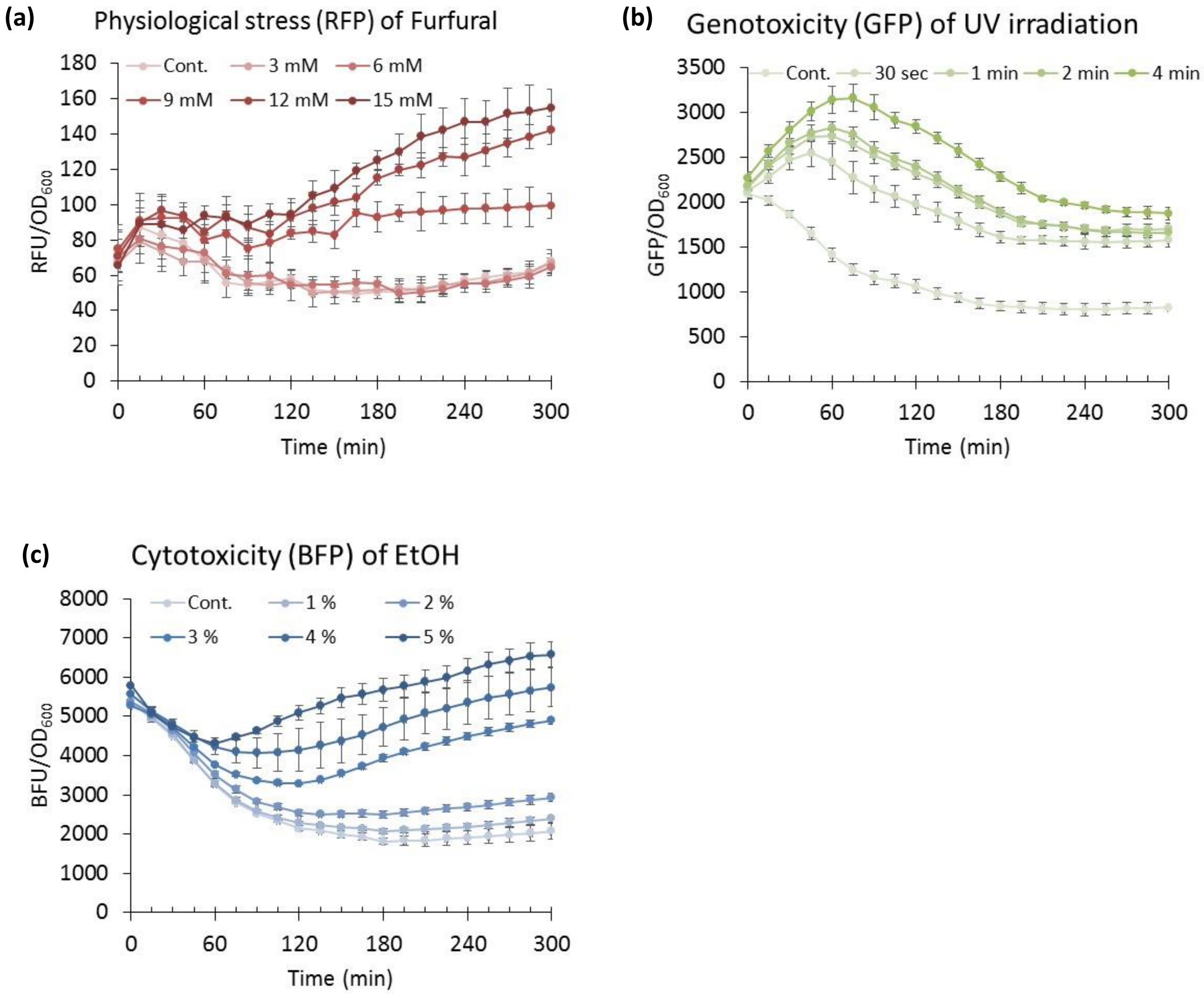


**Supplementary Figure 3: Kinetic analyses of stress responses.** The specific fluorescence of **(a)** RFP (physiological stress), **(b)** GFP (genotoxicity) and **(c)** BFP (cytotoxicity) induced by variable dosages of furfural, ultraviolet (UV) irradiation and ethanol, respectively, determined over 5 hours. Error bars are SD of six independent biological replicates. “Cont.” indicates negative control samples that were not treated with stressors.

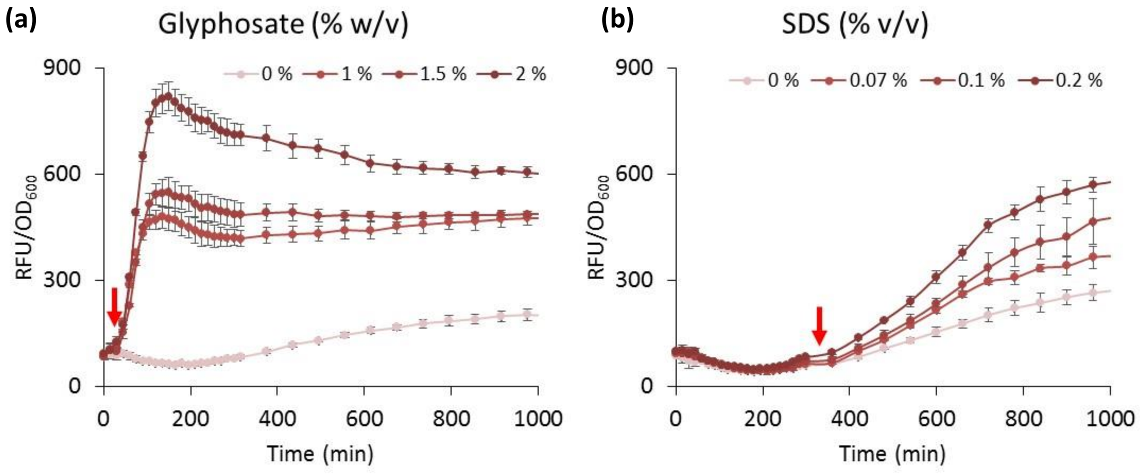


**Supplementary Figure 4**: **Variation in stress signal initiation times.** Variable physiological stress (RFP) signal initiation is observed in response to **(a)** glyphosate and **(b)** sodium dodecyl sulphate (SDS). Red arrows indicate times when the RFP signals discriminated from the control. Note the very quick response to glyphosate as compared to SDS. Error bars are the standard deviation (SD) of six biological replicates.

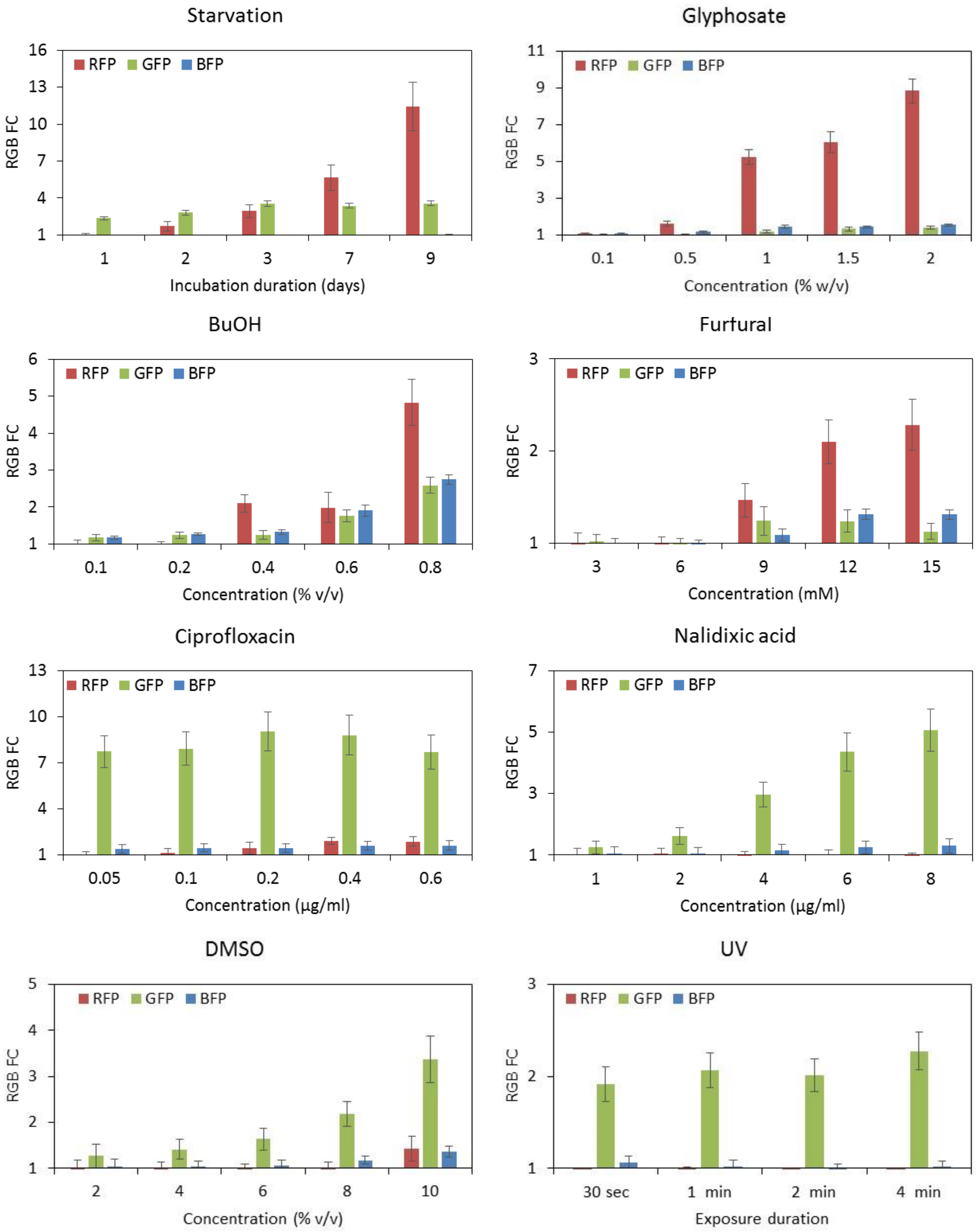
Starvation
RGB FC
RFP GFP BFP
1 4 7 10 13 16
1 2 3 7 9
Incubation duration (days)
Glyphosate
1 3 5 7 9 11
0.1 0.5 1 1.5 2
Concentration (% w/v)
BuOH
1 2 3 4 5 6
0.1 0.2 0.4 0.6 0.8
Concentration (% v/v)
Furfural
1 2 3
3 6 9 12 15
Concentration (mM)
Ciprofloxacin
1 4 7 10 13
0.05 0.1 0.2 0.4 0.6
Concentration (μg/ml)
Nalidixic acid
1 3 5 7
1 2 4 6 8
Concentration (μg/ml)
DMSO
1 2 3 4 5
2 4 6 8 10
Concentration (% v/v)
UV
1 2 3
30 sec 1 min 2 min 4 min
Exposure duration

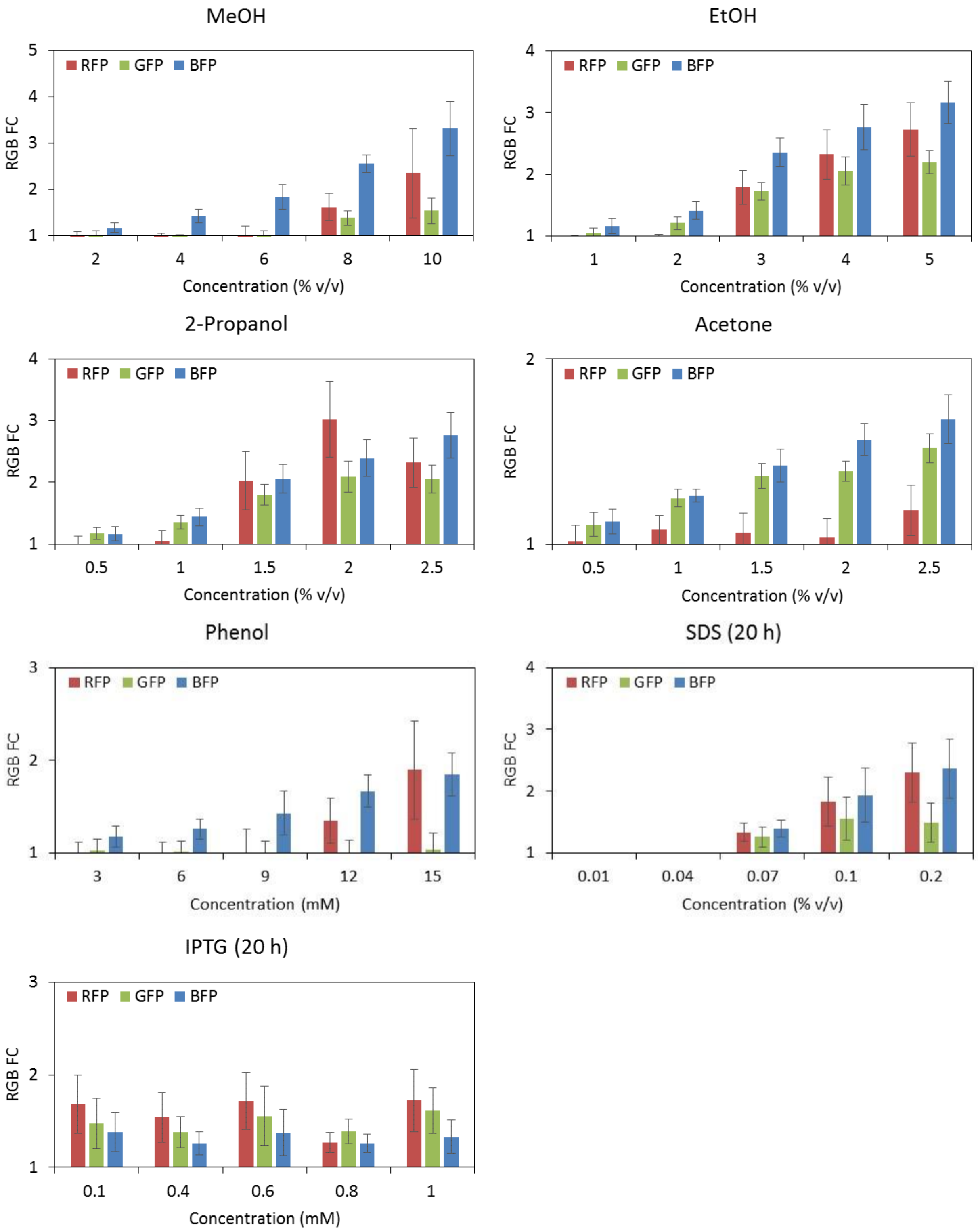


**Supplementary Figure 5: Dose dependent stress responses after exposure to various stressors.** The three fluorescence channels are represented as the fold change (RGB FC) of specific fluorescence ($FU/OD_{600}$) over the control culture. All cultures were incubated for 5 h in LB+kan medium, except SDS and IPTG were incubated for 20 h, and starvation was performed for 9 days. GFP is genotoxicity, RFP is physiological stress and BFP is cytotoxicity. Error bars are the standard deviation (SD) of six biological replicates.

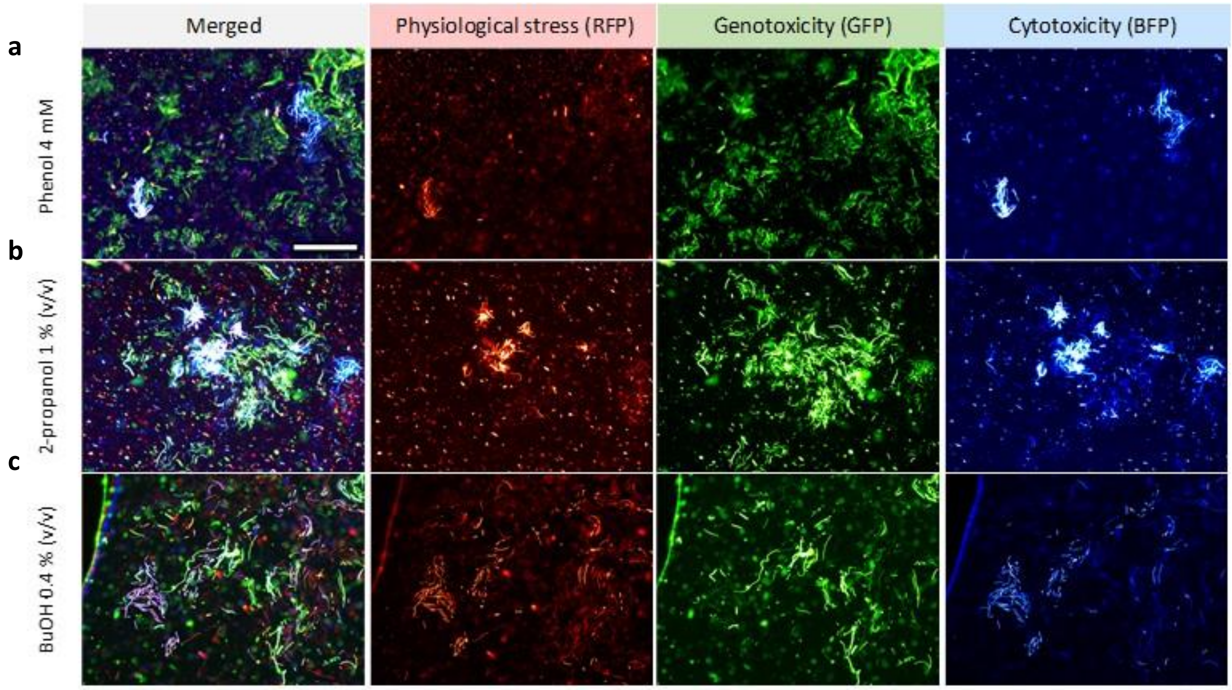


**Supplementary Figure 6: Phenotypic heterogeneity of stress responses in clonal populations.** Diverse subpopulations can be observed in response to phenol, 2-propanol and butanol after 24 h of planktonic growth. The three fluorescence channels for every stressor were imaged in the same field of view using identical instrument settings. Scale bar is 100 µm.

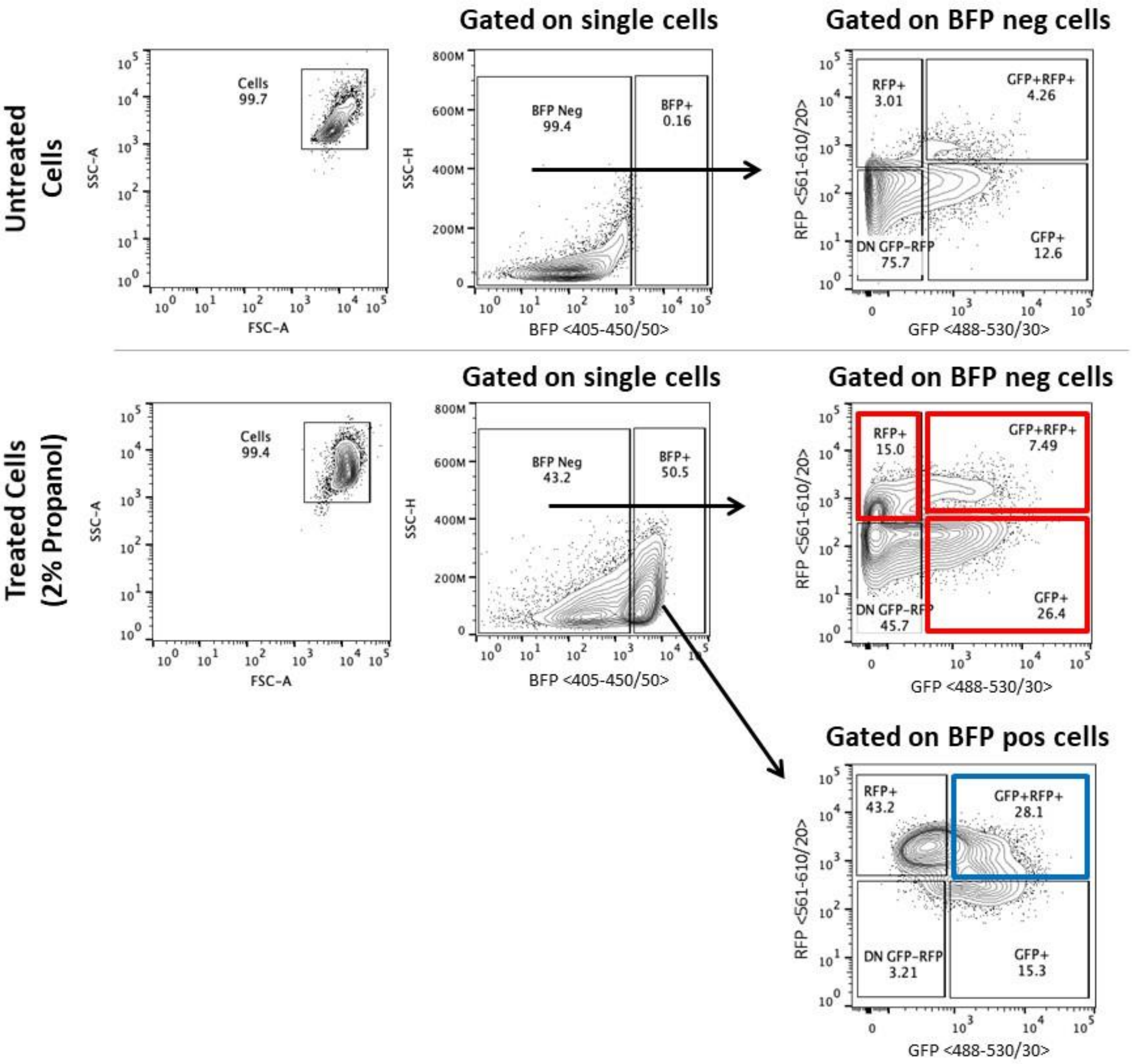


**Supplementary Figure 7: FACS profile of RGB-S *E. coli* treated cells. RGB-S *E. coli*** treated with 2-propanol 2% (v/v) isolated by FACS based on the single cell responses into four subpopulations; mono GFP (GFP+), mono RFP (RFP+), dual RFP-GFP (GFP+RPF+), and triple RFP-GFP-BFP (GFP+RPF+BFP+). Neg = negative, and pos = positive. Frequency of each population is depicted in the corresponding gate. Sorted populations are shown in red and blue gates. Representative figure of two replicates.

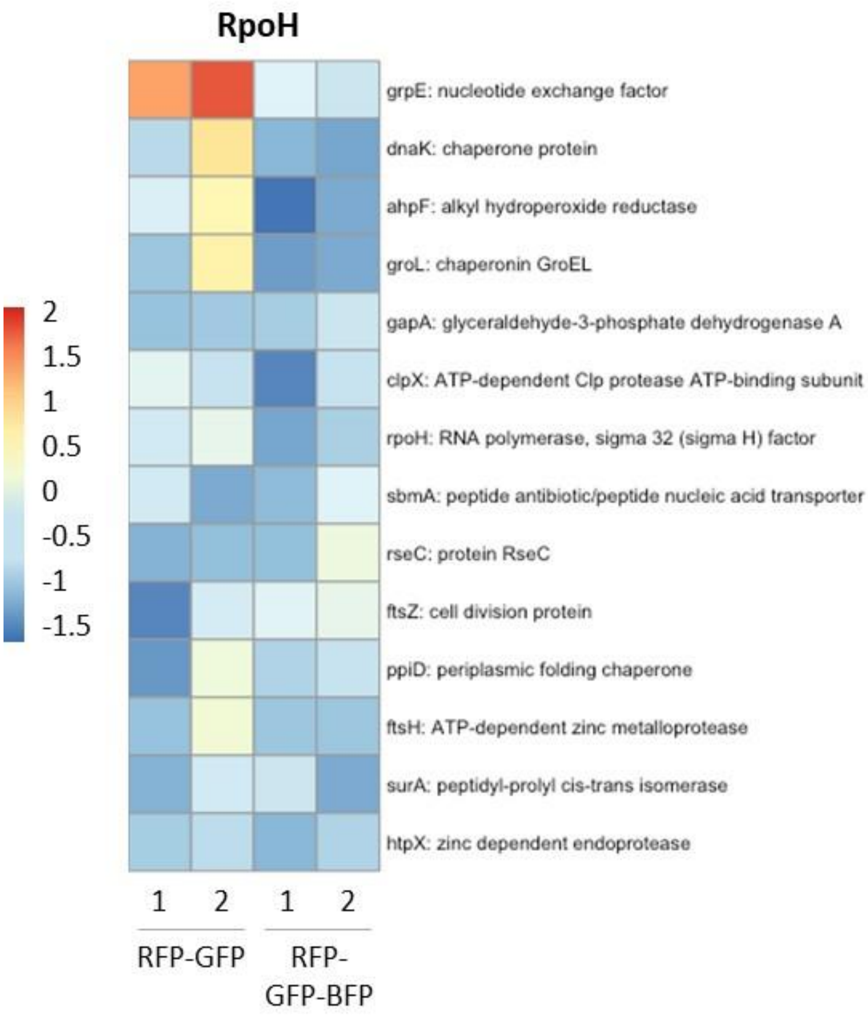


**Supplementary Figure 8: Transcriptomic analysis of RpoH responses.** RGB-S *E. coli* was treated with 2-propanol 2 % (v/v) and sorted using FACS based on the single cell responses into two subpopulations, dual RFP-GFP and triple RFP-GFP-BFP, with numbers (1, 2) indicating replicates. Differential transcription of genes involved in RpoH stress response pathway were analysed within RFP-GFP and RFP-GFP-BFP sorted subpopulations. The observed generally low transcription of the RpoH regulon in the triple response cells compared to the dual response suggests that those cells have largely shut down transcription, while still having accumulated all three fluorescent reporter proteins beforehand and thus being sorted as RGB positive.

(a)

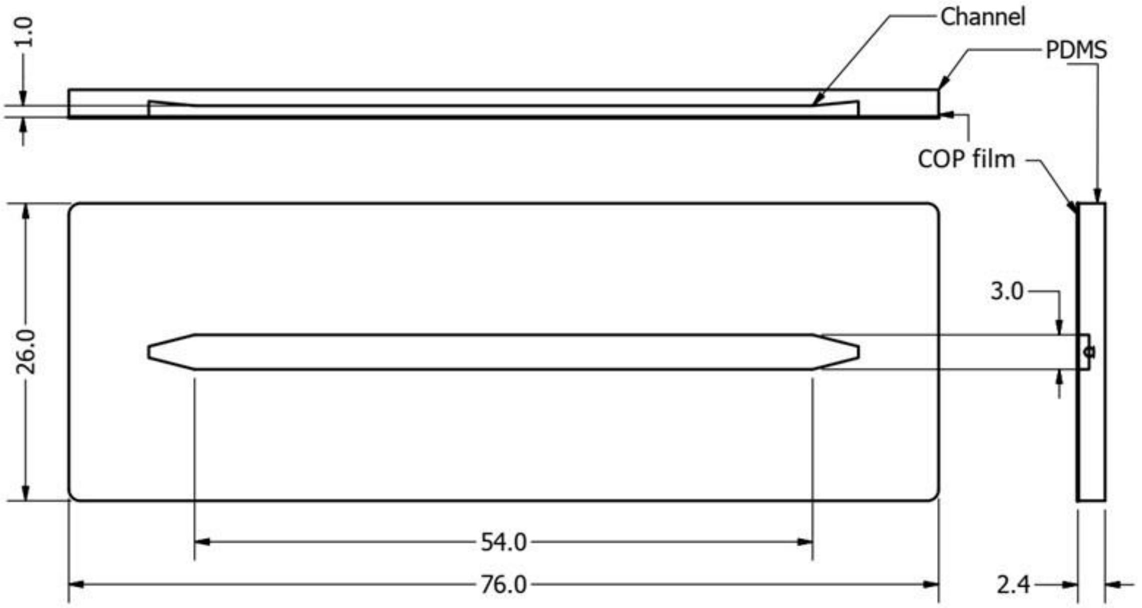


(b)

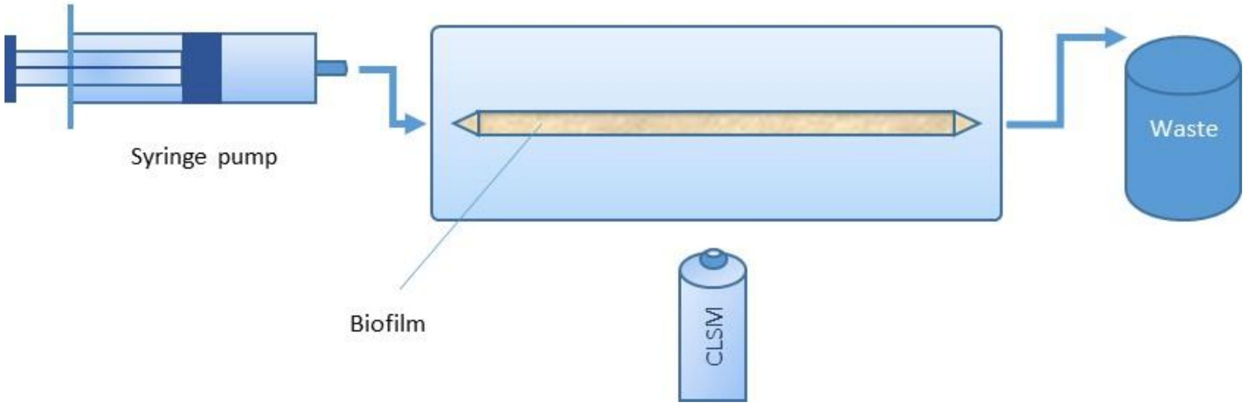


**Supplementary Figure 9: Setup of the microfluidic flow cell used for cultivation and 3D imaging of RGB-S *E. coli* biofilms. (a)** Technical illustration of the straight flow-cell microfluidic chip used to cultivate RGB-S *E. coli* biofilm. The chip is made of PDMS and bonded to a Cyclic Olefin Polymer (COP) film on which the biofilm was grown. All dimensions are in mm. **(b)** Setup of cultivating and imaging the stressed biofilm in the flow-cell chip using the confocal laser scanning microscope (CLSM).

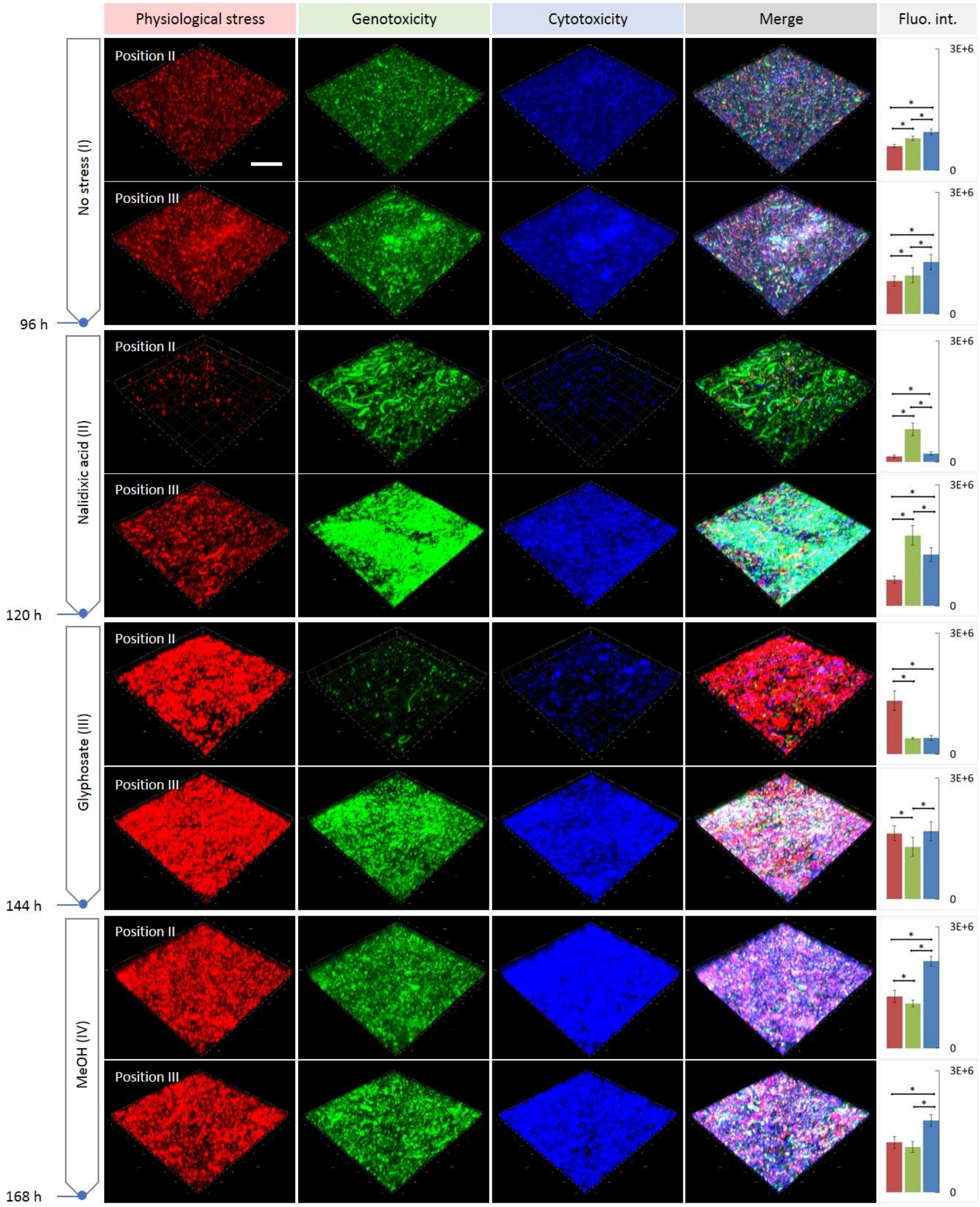


**Supplementary Figure 10: Responsiveness of *E. coli* biofilms to sequential stresses. (a)** Representative images taken at additional centre positions within the fluidic channel. Note that total fluorescence intensities are varying for the different positions due to changes in biomass distribution. The mature biofilm grown in the microfluidic chip under continuous LB+kan medium flow for 96 h mainly shows a basal level of stress response (no stress, **I**). The biofilm was sequentially treated with nalidixic acid (NA, **II**), glyphosate (Gly, **III**) and methanol (MeOH, **IV**), each for one day. This induced mainly the expected fluorescent protein expression; GFP (genotoxicity), RFP (physiological stress) and BFP (cytotoxicity), respectively. Scale bar is 100 µm. Quantitative red, green and blue bars (on the right) represent the intensity average of RFP, GFP and BFP, respectively, acquired from nine equal 100 $\mu m^2$ regions. (*) indicates statistical significance of response means. Error bars indicate standard deviation obtained from the analysis of nine individual 100 $\mu m^2$ field-of-view sections.

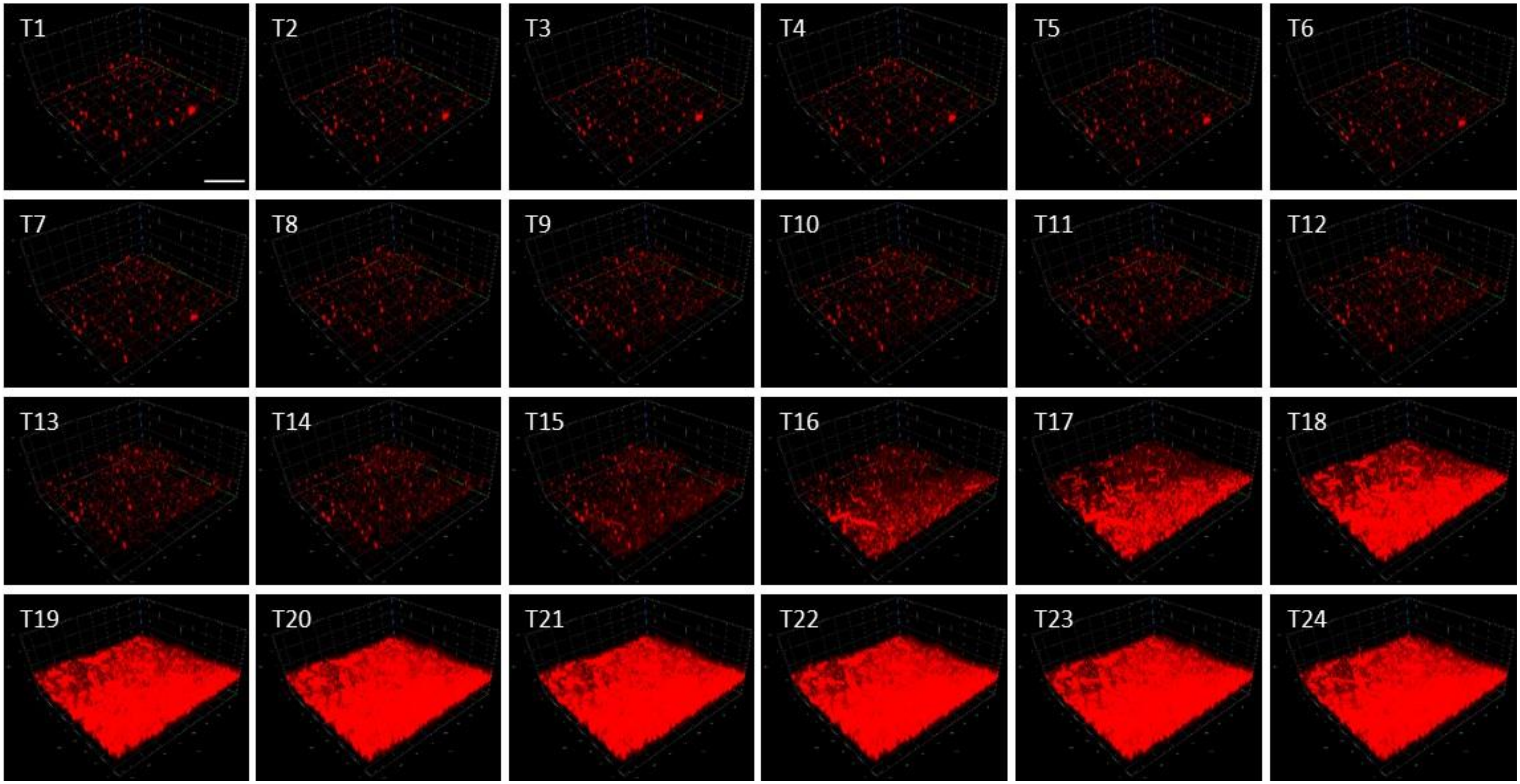


**Supplementary Figure 11: Real-time observation of the development of physiological stress response of RGB-S *E. coli* biofilm.** While exposing the biofilm to glyphosate 2 % (w/v) for 24 hours, an image was recorded at a centre position every 1 h (time points T1 - T24). Note that glyphosate administration induced a progressive RFP signal indicating physiological stress. Scale bar is 100 µm.

**Supplementary Table 1:** Wavelengths used for measurement of the three fluorescent proteins. Reference values and wavelength sets used for fluorescence measurements and image acquisition by instruments used in this study. A Synergy H1 (BioTek inc.) was used for quantitative readouts of the bulk cultures. A Zeiss fluorescence microscope was used for 2D imaging and a Zeiss CLSM was used for biofilm 3D imaging.

| **Fluorescent protein** | **Colour** | **Maxima (nm)** | **Synergy H1 filters (nm)** | **Fluorescence microscopy filters (nm)** | **CLSM filters (nm)** |
|---|---|---|---|---|---|
| mTagBFP2[1] | Blue | Ex: 399<br>Em: 454 | Ex: 400<br>Em: 454 | (Set 49) Ex: 365<br>Em: 445/50 | Ex: 405<br>Em: 410-530 |
| GFPmut3b[2] | Green | Ex: 501<br>Em: 511 | Ex: 483<br>Em: 511 | (Set 44) Ex: 475/40<br>Em: 530/50 | Ex: 488<br>Em: 490-597 |
| mRFP1[3] | Red | Ex: 584<br>Em: 607 | Ex: 571<br>Em: 607 | (Set 43 HE) Ex: 550/25<br>Em: 605/70 | Ex: 561<br>Em: 582-754 |

**Supplementary Table 2:** List of DNA sequences

| Part | Sequence |
|---|---|
| **Promoter *sulA* (SOS)** | TAGGGTTGATCTTTGTTGTCACTGGATGTACTGTACATCCATACAGTAACTCACAGGGGCTGGATTGATT |
| **Promoter *osmY* (RpoS)** | CTGGCACAGGAACGTTATCCGGACGTTCAGTTCCACCAGACCCGCGAGCATTAATTCTTGCCTCCAGGGCGCGGTAGCCGCTGCGCCCTGTCAATTTCCCTTCCTTATTAGCCGCTTACGGAATGTTCTTAAAACATTCACTTTTGCTTATGTTTTCGCTGATATCCCGAGCGGTTTCAAAATTGTGATCTATATTTAACAAAGTGATGACATTTCTGACGGCGTTAAATACCGTTCAATGCGTAGATATCAGTATCTAAAGCCGTCGATTGTCATTCTACCGATATTAATAACTGATTCAGAGGCTGTAATGGTCGTTATTCATCACTCATCGCTTTTGTGATGGCGACCATTGACTTCTGTAGAGGGTGAAGTCTCTCCCTATTCAGCAATGCAACCTCGTGTTGCCAGGCTCAAATTACGAGCAAACATACAGGAATAAATCG |
| **Promoter *grpE* (RpoH)** | GAATTTCTCCGCGTTTTTTTCGCATTCATCTCGCTAACTTCGCTTATTATGGGGATCAGTTTCAGGGTTTCAAGGGAAGCACTCACATTGTCATCAATCTTCGCAACAAGGACCTCGG |
| **Terminator_1 (double terminator)** | AGCCAGGCATCAAATAAAACGAAAGGCTCAGTCGAAAGACTGGGCCTTTCGTTTTATCTGTTGTTTGTCGGTGAACGCTCTCTACTAGAGTCACACTGGCTCACCTTCGGGTGGGCCTTTCTGCGTTTATATACTAGA |
| **Terminator_2 (double terminator)** | TACTAGAGCCAGGCATCAAATAAAACGAAAGGCTCAGTCGAAAGACTGGGCCTTTCGTTTTATCTGTTGTTTGTCGGTGAACGCTCTCTACTAGAGTCACACTGGCTCACCTTCGGGTGGGCCTTTCTGCGTTTATA |
| **Terminator_3 (double terminator)** | CTCGGTACCAAATTCCAGAAAAGAGGCCTCCCGAAAGGGGGGCCTTTTTTCGTTTTGGTCCCCAATTATTGAAGGCCGCTAACGCGGCCTTTTTTTGTTTCTGGTCTCCC |
| **Terminator_4 (single terminator)** | CAGATAAAAAAAATCCTTAGCTTTCGCTAAGGATGATTTCT |
| ***mRFP1* (RFP, codon optimized)** | ATGGCAAGCAGCGAAGATGTGATCAAAGAATTTATGCGTTTCAAGGTGCGTATGGAAGGTAGCGTTAATGGTCATGAATTTGAAATCGAAGGTGAAGGTGAGGGTCGTCCGTATGAAGGCACCCAGACCGCCAAACTGAAAGTTACCAAAGGCGGTCCGCTGCCGTTTGCATGGGATATTCTGAGTCCGCAGTTTCAGTATGGTAGCAAAGCATATGTTAAACATCCGGCAGATATCCCGGATTATCTGAAACTGAGCTTTCCGGAAGGCTTTAAATGGGAACGTGTGATGAATTTTGAGGATGGTGGTGTTGTTACCGTTACACAGGATAGCAGCCTGCAGGATGGTGAATTTATCTATAAAGTTAAACTGCGTGGCACGAATTTTCCGTCAGATGGTCCTGTTATGCAGAAAAAAACCATGGGTTGGGAAGCAAGCACCGAACGTATGTATCCGGAAGATGGTGCACTGAAAGGTGAAATCAAAATGCGTCTGAAGCTGAAAGATGGCGGTCATTATGATGCAGAAGTTAAAACCACCTACATGGCCAAAAAACCGGTGCAGCTGCCTGGTGCATATAAAACGGATATTAAACTGGATATCACCTCGCACAACGAGGATTATACCATTGTTGAACAGTATGAACGTGCAGAAGGTCGTCATAGTACCGGTGCCTAATAA |
| ***GFPmut3b* (GFP, codon optimized)** | ATGCGTAAAGGTGAAGAACTGTTTACCGGTGTTGTTCCGATTCTGGTTGAACTGGATGGTGATGTTAATGGCCACAAATTTTCAGTTAGCGGTGAAGGCGAAGGTGATGCAACCTATGGTAAACTGACCCTGAAATTTATCTGTACCACCGGCAAACTGCCGGTTCCGTGGCCGACACTGGTTACCACCTTTGGTTATGGTGTTCAGTGTTTTGCACGTTATCCGGATCATATGAAACAGCACGATTTTTTCAAAAGCGCAATGCCGGAAGGTTATGTTCAAGAACGTACCATCTTCTTCAAAGATGACGGCAACTATAAAACCCGTGCCGAAGTTAAATTTGAAGGTGATACCCTGGTGAATCGCATTGAACTGAAAGGCATCGATTTTAAAGAGGATGGTAATATCCTGGGCCACAAACTGGAATATAATTATAATAGCCACAACGTGTACATCATGGCCGACAAACAGAAAAATGGCATCAAAGTGAACTTCAAGATCCGCCATAAT |

| | |
|---|---|
| | ATTGAAGATGGTTCAGTTCAGCTGGCCGATCATTATCAGCAGAATACCCCGATTGGT GATGGTCCGGTTCTGCTGCCGGATAATCATTATCTGAGCACCCAGAGCGCACTGAG CAAAGATCCGAATGAAAAACGTGATCACATGGTGCTGCTGGAATTTGTTACCGCAG CAGGTATTACCCATGGTATGGATGAACTGTATAAATGATAA |
| ***mTagBFP2* (BFP, codon optimized)** | TTATCAGTTCAGTTTATGACCCAGTTTGCTCGGCAGATCACAATAACGTGCAACTGC AACTTCATGCTGTTCAACATAGGTTTCGTTGTTTGCTTCTTTGATGCGTTCCAGACGA TAATCCACATAATAAACGCCAGGCATTTTCAGATTCTTTGCCGGTTTTTTGCTACGAT AGGTGGTTTTGGCATTTGCAATCAGATGGCTACCACCAACCAGTTTCAGTGCCATAT CATTACGACCTTCCAGGCCACCATCTGCCGGATACAGTGTTTCGGTAAATGCTTCCC AACCTAAGGTCTTTTTCTGCATAACCGGACCATTGCTGGTGAAGTTCACACCGCGAA TTTTCACATTATAAATCAGACAACCATCCTGCAGACTGGTATCCTGTGTTGCGGTCAG AACACCACCATCTTCATAGGTGGTAACACGTTCCCAGGTAAAACCTTCCGGAAAGCT CTGTTTGAAAAAATCCGGGATACCCTGGGTATGATTGATAAAGGTTTTGCTACCATA CAGAAAGCTGGTTGCCAGAATATCAAATGCAAACGGCAGCGGACCACCTTCAACAA CTTTAATACGCATGGTCTGGGTGCCTTCATACGGTTTACCTTCACCTTCGCTGGTACA TTTAAAGTGGTGGTTATCAACGGTGCCTTCCATATACAGTTTCATGTGCATGTTTTCT TTGATCAGTTCTTCGCCTTTGCTAACCAT |
| **O17051-F primer** | CCAGTGAGCGCGACGTAATA |
| **O17052-R primer** | CGGCCACTCAACCCTATCTC |
